# Imaging cell lineage with a synthetic digital recording system

**DOI:** 10.1101/2020.02.21.958678

**Authors:** Ke-Huan K. Chow, Mark W. Budde, Alejandro A. Granados, Maria Cabrera, Shinae Yoon, Soomin Cho, Ting-hao Huang, Noushin Koulena, Kirsten L. Frieda, Long Cai, Carlos Lois, Michael B. Elowitz

## Abstract

Multicellular development depends on the differentiation of cells into specific fates with precise spatial organization. Lineage history plays a pivotal role in cell fate decisions, but is inaccessible in most contexts. Engineering cells to actively record lineage information in a format readable *in situ* would provide a spatially resolved view of lineage in diverse developmental processes. Here, we introduce a serine integrase-based recording system that allows *in situ* readout, and demonstrate its ability to reconstruct lineage relationships in cultured stem cells and flies. The system, termed intMEMOIR, employs an array of independent three-state genetic memory elements that can recombine stochastically and irreversibly, allowing up to 59,049 distinct digital states. intMEMOIR accurately reconstructed lineage trees in stem cells and enabled simultaneous analysis of single cell clonal history, spatial position, and gene expression in *Drosophila* brain sections. These results establish a foundation for microscopy-readable clonal analysis and recording in diverse systems.

**One sentence summary:** A new genetic editing system termed intMEMOIR reveals the lineage histories of individual cells directly within their native tissue context.

## Main Text

Cell lineage plays pivotal roles in cell fate determination in development, homeostasis, and disease (*1*–*5*). The ability to visualize lineage relationships directly within their native tissue context would provide powerful insights into the roles of intrinsic and extrinsic factors in cell fate specification. Recently, an explosion of engineered lineage recording systems have promised to revolutionize lineage analysis. Inspired by the recovery of lineage information from naturally occurring somatic mutations (*2*, *3*, *5*–*12*), these systems actively generate stochastic, heritable mutations at defined genomic target sites, and then recover those edits in individual cells to reconstruct their lineage (*3*, *11*, *13*–*23*). Currently, most of these methods require readout by sequencing, which disrupts spatial organization entirely. Another approach, Memory by Engineered Mutagenesis with Optical In situ Readout (MEMOIR) uses single-molecule fluorescence *in situ* hybridization (smFISH) to allow readout by imaging. However, it relies on genomically distributed deletion edits that do not permit extended recording and germline transmission (*23*). Thus, a powerful, broadly useful image-readable recording system has been lacking.

To address this challenge, we developed a digital, image-readable lineage recording system termed intMEMOIR. The system introduces a novel design based on a dense multi-state memory array that can be edited by serine integrases and integrated at defined genomic sites for germline heritability, allowing its use in diverse organisms and contexts. This design generates diverse, digital, irreversible edits states that can be stably inherited over many cell cycles and then read out by smFISH alongside endogenous transcripts (*24*–*26*). We show that intMEMOIR accurately reconstructs lineage information in mouse embryonic stem cells, and links lineage, cell fate, and spatial structure in the adult *Drosophila* brain.

## Results

### Serine integrases enable FISH-readable three-state memory elements

One mode of lineage reconstruction is clonal tracing, which identifies cells descended from a common ancestor based on unique, heritable labels (e.g. sequence barcodes). A more complete lineage reconstruction further provides the tree, or pedigree, of multiple divisions through which cells are related (Fig. 1A). To access both regimes, a recording system should be able to produce and preserve as much molecular diversity as possible to maximize the number of distinguishable clones, and accumulate that diversity over multiple cell generations to enable tree reconstruction. In systems based on two-state memory elements (bits), extended recording durations eventually edit all memory elements, producing a non-informative homogeneous end state, and effectively erasing recorded information. By contrast, three-state memory elements, or trits, that start in an initial state and irreversibly switch to one of two potential end states, provide additional information per element and preserve recorded information. As a result, the use of trits dramatically improves the accuracy of multi-generation reconstruction and the multiplexibility of clonal classification in simulations, and allows the system to function across a broader range of edit rates compared to bit-based memory (Fig. 1B and C) (*27*).

**Fig. 1.**
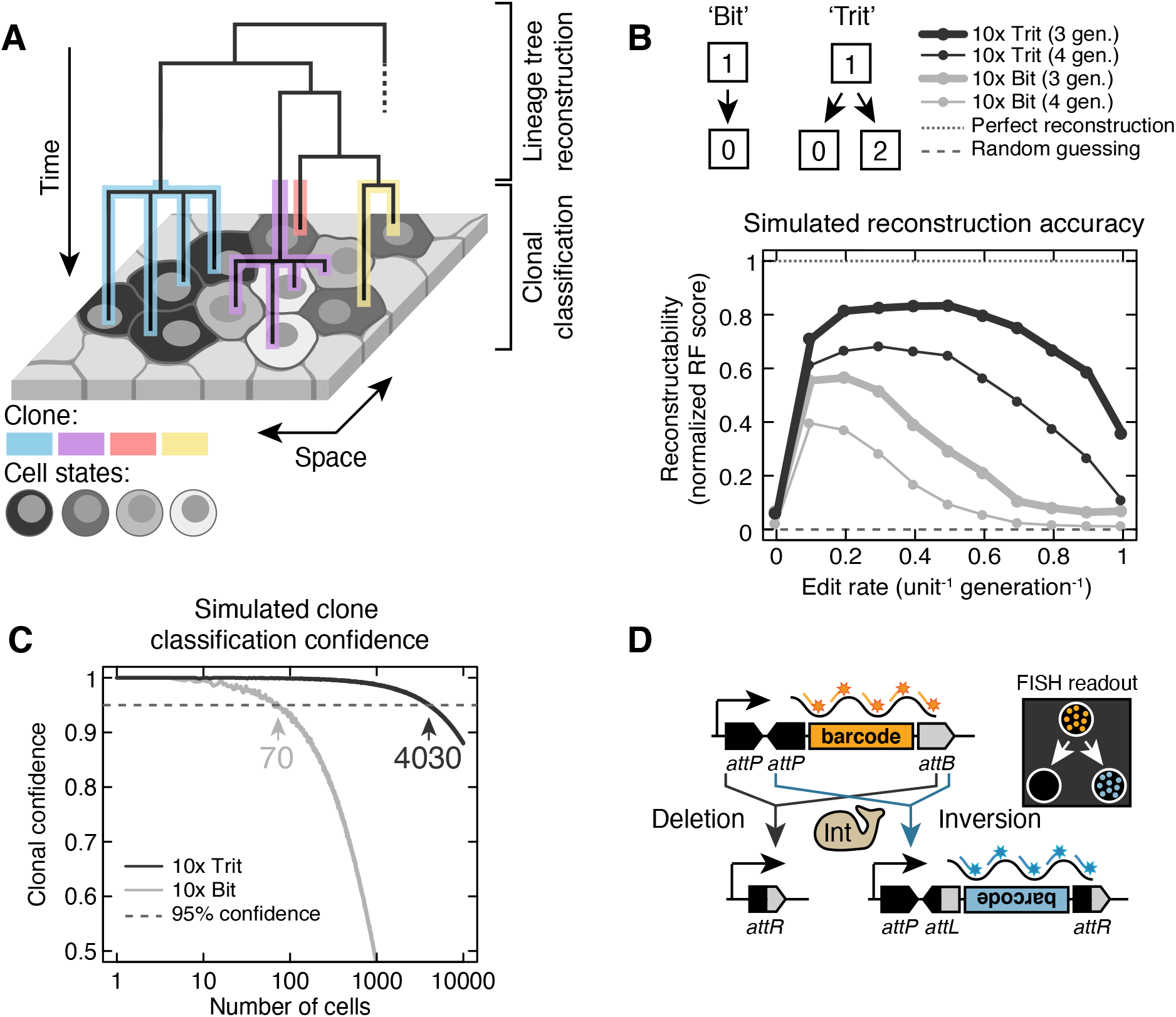
Three-state memory elements (trits) enable *in situ* developmental lineage reconstruction. (**A**) An ideal recording system connects the spatial information, gene expression, and lineage history of single cells (schematic). Lineage information comprises multi-generation reconstruction of cell division trees (panel B) as well as multiplexed clone classification (panel C). (**B** and **C**) Simulations of editing and reconstruction for systems with 10 irreversible recording units. For lineage tree reconstruction in (B), trit recording units improve tree reconstructability compared to two-state bits across a wide range of edit rates, and retain information when edited to completion. Reconstructability is defined as the normalized Robinson-Foulds score obtained by comparing the reconstructed tree to ground truth simulated lineage. For clonal classification in (C), trits enable simultaneous tracing of a large number of clones in the same organism, potentially distinguishing up to two orders of magnitude more clones with >95% confidence than its bit counterpart (4030 and 70 clones, respectively). Clonal confidence is defined as the probability that two randomly selected cells with identical edit patterns are from the same clone. (**D**) Serine integrases enable trit designs compatible with FISH readout. Transcribed barcodes are flanked by two *attP*s and one *attB* site. Recombination results in either an inverted or deleted barcode, which can be distinguished by fluorescent probes (colored lines and asterisks) directed against either strand.

Phage serine integrases provide an ideal basis for engineering trits. They mediate irreversible recombination between directional *attP* and *attB* target sites, deleting or inverting the intervening sequence depending on relative site orientation (*28*–*32*). Serine integrases such as Bxb1 do not rely on endogenous repair mechanisms to generate edits, and they function across species, including in mammalian cells (*32*–*34*). To create a trit, we flanked a barcode sequence by an inverted pair of *attP* sites on one end and an *attB* site on the other such that Bxb1-mediated recombination produces either irreversible barcode deletion or inversion (Fig. 1D). A strong Pol II promoter drives transcription of the trit, allowing *in situ* readout by smFISH. Before editing, the promoter expresses the forward barcode, whereas after recombination, it expresses either no transcript (deletion) or the reverse complement barcode (inversion), enabling digital discrimination of trit states by smFISH.

Concatenating multiple trits in a compact array and integrating it at a safe harbor locus (*35*–*37*) increases the amount of memory while facilitating germline transmissibility (*38*). Edit strategies that rely on double-stranded breaks and endogenous DNA repair machinery are prone to information loss in arrays through inter-unit deletions (*27*, *39*). By contrast, integrases allow recombinational isolation between distinct trits in the same array. Within each *att* site, a central dinucleotide confers both specificity and directionality of recombination (*34*, *40*) (Fig. 2A). In principle, 10 distinct dinucleotide variants can be used orthogonally (Fig. S1), enabling 10 corresponding independent memory units in a single array, for a theoretical diversity of 3^10^ (59,049) states.

**Fig. 2.**
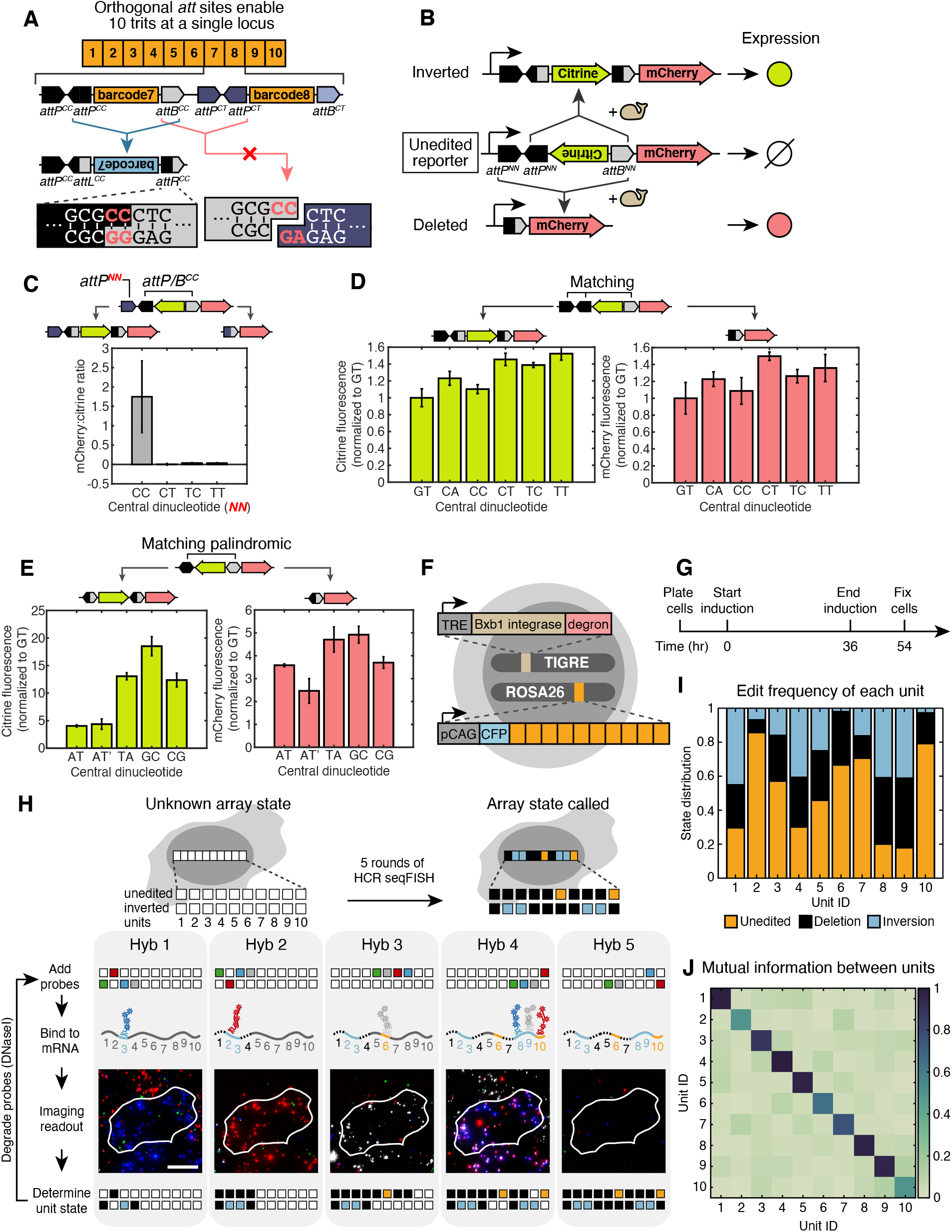
Trits can be independently edited within recording arrays. (**A**) 10 trits (orange numbered rectangles) can be concatenated into a 10-unit array using *attP*/*attB* pairs with orthogonal central dinucleotides (red letters). (**B**) Fluorescent reporter assay enables rapid characterization of the recording units. The trit reporter construct (middle) produces no fluorescence until an integrase inverts it to express Citrine (top) or deletes it to express mCherry (bottom). In subsequent panels, cells transfected with reporter constructs were analyzed by flow cytometry, gating on a co-transfection marker (Materials and Methods). Data in subsequent panels are calculated from median values of triplicate experiments. (**C**) Bxb1 does not efficiently recombine *attP* and *attB* with mismatched dinucleotides. (**D**) All six non-palindromic dinucleotides mediate inversion (left) and deletion (right) of *attP/B* pairs. (**E**) With palindromic dinucleotides, one pair of matching *attP* and *attB* is sufficient for inversion and deletion. Bxb1 is agnostic to the relative orientation of these *att* sites, as demonstrated by the comparable edit efficiencies between *attP/B*^AT^ when the two sites are arranged in the same (AT) and opposite (AT’) orientations. (**F**) intMEM1 is a stable mES cell line with the 10-unit array integrated at the Rosa26 locus and an inducible Bxb1 integrated at the TIGRE locus. The 10-unit array is constitutively expressed, while Bxb1 can be activated by the combination of doxycycline for transcription, and trimethoprim (TMP) for protein stabilization. (**G**) Over 36 hours of growth with Bxb1 induction, cells progressively accumulate edits for multi-generational lineage reconstruction. Induction is then stopped, and array edits are inherited by daughter cells over an 18 hour expansion period, enabling clonal classification. (**H**) Five rounds of hybridization read out all possible states of the recording array *in situ*. (**I**) The relative frequency with which each trit is observed in its unedited, deleted, or inverted state after 36 hours of Bxb1 induction. (**J**) The low mutual information between any given pair of units illustrates functional independence of each of the 10 units in the array.

To validate the trit design in mouse embryonic stem (mES) cells, we constructed a prototype Bxb1 trit that expressed no fluorescent protein, Citrine, or mCherry in the unedited, inverted, and deleted states, respectively, allowing rapid characterization by flow cytometry (*33*) (Fig. 2B). Recombination occurred much more efficiently between matching compared to mismatching dinucleotides, indicating that distinct dinucleotides operate in a largely orthogonal manner, as intended (Fig. 2C). When the *attB* and inverted *attPs* all contained the same dinucleotide, such as GT, inversion and deletion both occurred efficiently (Fig. 2D). Further, *att* sites made from palindromic dinucleotides such as AT, which lack directionality, enable a simplified design in which a single *attP!B* pair mediates both inversion and deletion (Fig. 2E). A control construct in which one site is inverted (AT’) performed similarly to the uninverted counterpart, permitting the use of palindromic sites in either orientation (Fig. 2E). This design could also be generalized to other integrases (Fig. S2). Although deletion crosstalk did occur at low rates (Fig. 2C), consistent with observations using phiC31 integrase (*41*), these results show that dinucleotide variants permit orthogonal recombination, enabling construction of compact 10-unit trit arrays.

### Design and characterization of the intMEM1 recording cell line

Based on these results, we engineered a cell line, intMEM1, capable of inducible autonomous recording. We constructed an array of 10 trits and site-specifically integrated it at the ROSA26 locus in mES cells (Fig. 2F). We also site-specifically integrated an inducible Bxb1 cassette at the TIGRE safe-harbor locus (*42*). In this construct, doxycycline controls Bxb1 transcription and trimethoprim (TMP) stabilizes the protein by inhibiting a fused ecDHFR degron sequence (*43*). These complementary, redundant control systems together ensure tight regulation of integrase activity.

To quantify editing rates and outcomes, we co-cultured a low density of intMEM1 cells together with an excess of unengineered parental cells to support their growth. We induced recording for 36 hours by addition of doxycycline and TMP, continued growth for 18 additional hours without inducers, and then fixed the cells with formaldehyde (Fig. 2G). We used five sequential rounds of hybridization chain reaction smFISH (HCR-FISH) (*44*) to read each trit’s edit state. Further, we subsequently imaged the same cells by immunofluorescence with antibodies against membrane proteins E-cadherin and p-catenin to facilitate segmentation of adjacent cells in images.

The state of the entire array can be determined with five rounds of imaging using four fluorophores and twenty unique probes, one for each orientation of each trit (Fig. 2H). For example, in Figure 2H, the first imaging round revealed a signal for the inverted orientation of trit 3 and no signal for the inverted orientations of trits 1 and 4, nor for the unedited orientation of trit 2. The second hybridization probed the opposite states of the same four trits, revealing inversion of trit 2, and deletion of trits 1 and 4. In this way, we determined the full array editing outcomes in 1,487 array-expressing cells. Most trits were deleted and inverted at similar rates (Fig. 2I). However, trits 6 and 10 underwent deletions but rarely inverted. DNA sequencing revealed that trit 10 lacked *attP* sites, but trit 6 sequence was intact. Most importantly, mutual information analysis showed that trits within the same array were edited independently (Fig. 2J). Taken together, these results demonstrate that trits can be combined into a compact recording array of independent units.

### intMEMOIR reconstructs lineage relationships

To quantify intMEMOIR’s lineage reconstruction ability, we obtained ground-truth lineages from time-lapse movies and compared them with lineage relationships reconstructed from array edits in the same cells (Fig. 3A). We induced Bxb1 expression for 36h (Fig. 2G) to achieve −3 generations of recording, followed by an additional −1-2 generations of clonal expansion without Bxb1 induction, resulting in colonies of 13.7±7.7 cells (mean±s.d.). We then read out the state of the array using smFISH, classified the cells into clones corresponding to unique edit patterns, and performed multi-generation lineage tree reconstruction.

**Fig. 3.**
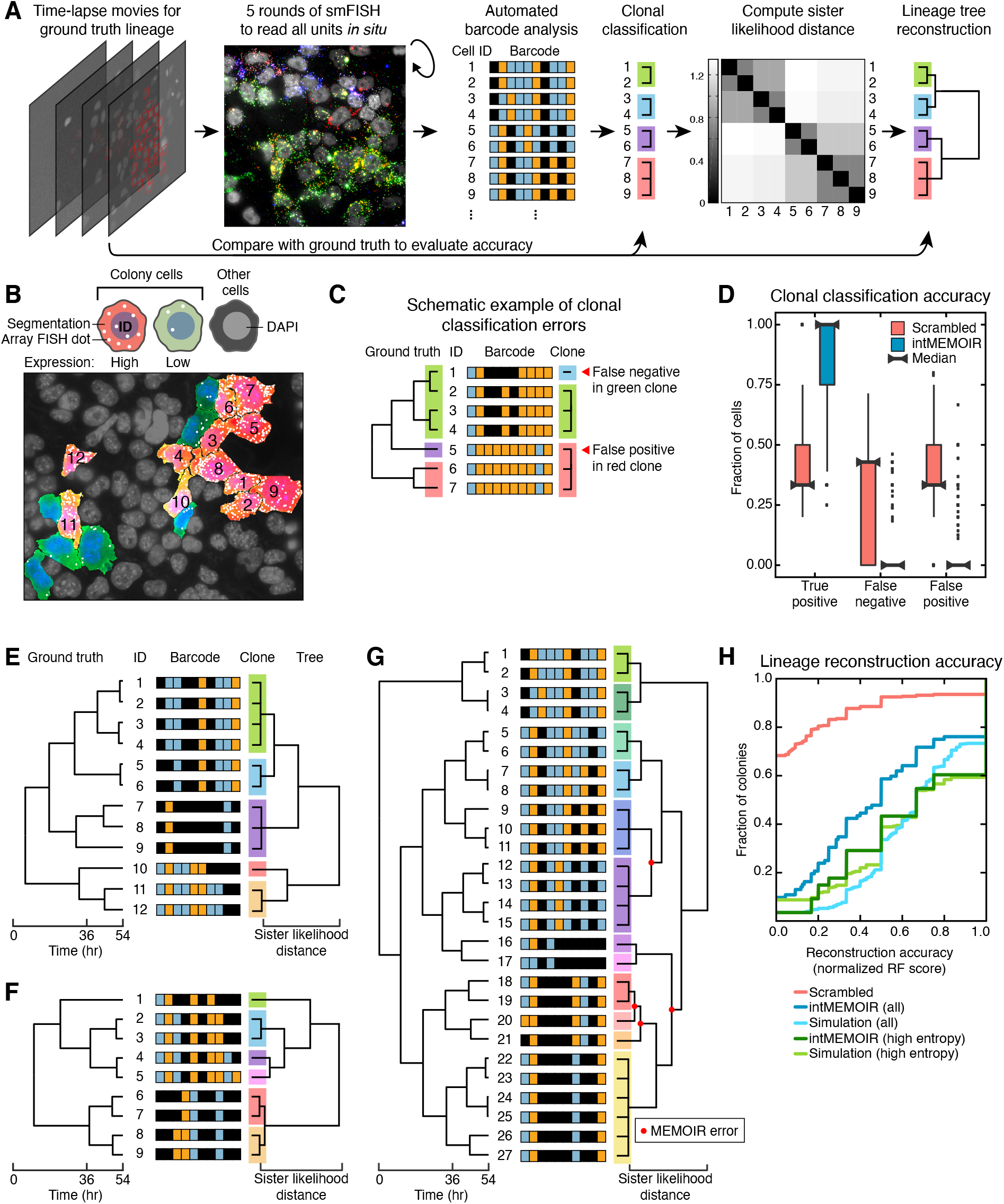
intMEMOIR reconstructs lineage relationships. (**A**) Quantitative analysis of the accuracy of clonal reconstruction and multi-generational lineage reconstruction. intMEM1 cells are tracked in time-lapse microscopy to establish ground truth lineage relationships (left panel). End-point FISH analysis recovers array edit states (second panel). Distinct edit patterns are used to classify cells into clones (‘clonal classification’, color groups), or further analyzed based on sister likelihood distance (Supplementary Text and Materials and Methods) to reconstruct lineage trees (‘lineage tree reconstruction’, right). Comparison with ground truth allows quantification of reconstruction accuracy. (**B**) Cells from the same colony are segmented (highlighted in red and green) and their array RNA FISH dots identified (white dots). Downstream analyses are performed on cells with strong array expression (red cells). (**C**) Clonal classification errors arise either from clonal cells gaining additional edits (false negative) or convergent edits between distant relatives (false positive). (**D**) intMEMOIR demonstrates robust clonal classification accuracy, with few errors compared to scrambled control. (**E-G**) Lineage reconstruction examples, with ground truth lineage on the left, cell ID and their corresponding barcode states in the middle, and reconstructed lineage on the right. (E) and (F) are colonies with perfect tree reconstruction, while (G) shows reconstruction error in branches highlighted (red dots). (**H**) Cumulative distributions show that lineage reconstruction from intMEMOIR approaches the accuracy expected from simulations of a 10-unit trit array displaying experimentally observed edit rates (blue, solid and dashed lines, respectively). Higher observed entropy in the edit patterns can independently identify colonies with greater reconstruction accuracy (green).

To assess clonal identification accuracy, we quantified the number of distinct edit states that could be detected in each colony, and the fidelity with which they reflected ground truth clonal relationships (Fig. S3 and Supplementary Text). We focused on the 76% of intMEM1 cells within colonies that showed the strongest array expression (Fig. 3B, orange vs. green cells), and omitted colonies with fewer than 3 states that are trivial to reconstruct. These colonies exhibited 8.4±4.7 (mean±s.d) unique array states. Across 1,453 cells spanning 105 colonies, 318 unique array edit patterns appeared in two or more cells in the same colony, with most clones (289) comprising 2-4 cells. Most clones were classified perfectly (median accuracy = 100%), and the average percentage of correctly classified cells per reconstructed clone was 85%, far exceeding results from a negative control analysis in which barcode-cell relationships were scrambled (Fig. 3D). Errors reflected false negative ambiguities due to subsets of cells within a clone undergoing additional edits, and false positive events in which distantly related cells convergently edited to identical patterns (Fig. 3C and Fig. S3). On average, these errors occurred in <10% of cells per clone, and more than half of the clones had an error rate of 0% (Fig. 3D). Thus, intMEMOIR performed accurate clonal classification.

Next, we assessed the ability to reconstruct lineage trees (Fig. 3A, ‘lineage tree reconstruction’). We used a maximum likelihood approach that incorporates the empirically determined recording parameters (Fig. 2I), assuming a constant edit rate per unedited site (Fig. S4 and Supplementary Text). Using this framework, we computed the probability of observing each array state after *G* generations starting from an unedited array; the conditional probability of observing any two specific array states as a pair of sister cells (see Supplementary Text); and, based on these probabilities, the relative likelihood of observing a given pair of array states for two sister cells compared to two unrelated cells. This likelihood provided a pairwise distance metric, which we then used to reconstruct a hierarchical lineage tree (Fig. 3, E to G and Table S2, see Supplementary Text). Reconstructed trees for the classified clones were often identical (Fig. 3E and F) or at least strikingly similar (Fig. 3G) to corresponding ground truth lineage trees. In some cases, reconstruction errors could be attributed to multi-trit deletions at the 3’ end of the array (e.g. Fig. 3G, cells 16 and 17).

To quantify reconstruction fidelity, we employed the Robinson-Foulds (RF) metric, defined as the fraction of lineage partitions (clades) that are shared between the reconstruction and the ground truth (*45*). We defined a normalized RF score (Supplementary Text) that ranges from 0 (complete disagreement) to 1 (perfect agreement). Stochastic simulations provided an upper bound to the possible accuracy of the system under ideal conditions, given the memory capacity of a single array, the empirically measured editing rates, and the observed set of ground truth trees (Fig. 3H, black). In parallel, we repeated this analysis, randomizing the cell-barcode relationships in the ground truth lineage, to compute a lower bound on reconstruction performance (Fig. 3H, red). Among the actual tree reconstruction scores (Fig. 3H, blue), 25% of colonies reconstructed perfectly (RF score = 1), while the overall score distribution was significantly higher than the random control (p <10^−16^, Kolmogorov-Smirnov test). Further, colonies with greater normalized entropy in their edit patterns reconstructed with higher accuracy (Fig. S5). For instance, trees with the 40% highest normalized entropy showed performance similar to that of simulated optimal recording, statistically equivalent to the upper bound (p>0.2, K-S test) (Fig. 3H, green). In applications where no ground truth is available, the entropy score thus allows one to enrich for subsets of cells likely to reconstruct with greater accuracy. Together, these results indicate that lineage recording and reconstruction can approach theoretical limits. The use of two or more arrays should therefore reconstruct colonies with greater depth and accuracy, as shown through simulations (Fig. S6) (*27*, *46*).

### intMEMOIR reconstructs early lineage of large mES colonies

In many developmental contexts, it is of interest to know how distinct clones that acquired different fates were related to each other at an earlier time point (*2*, *47*). As a proof of principle for this type of analysis, we followed 36h of editing with an additional 70h of growth without editing (−6 cell divisions). We then fixed cells, analyzed array states, and classified clones (Fig. 4A). Imaging revealed large domains of distinct, non-redundant edits in each array element (Fig. 4B). Combining these images provided a spatial map of clonal boundaries (Fig. 4C). Clonality broadly correlated with spatial position, as expected for colony growth. However, all clones were spatially extended and non-contiguous (Fig. 4C). Thus, intMEMOIR’s ability to generate high digital diversity enables it to simultaneously label many intermingled clones. Further, the specific edit patterns also allow inference of clonal relatedness (Fig. 4D, right). Together, these results show how intMEMOIR can be used as a powerful image-based clonal mapping system.

**Fig. 4.**
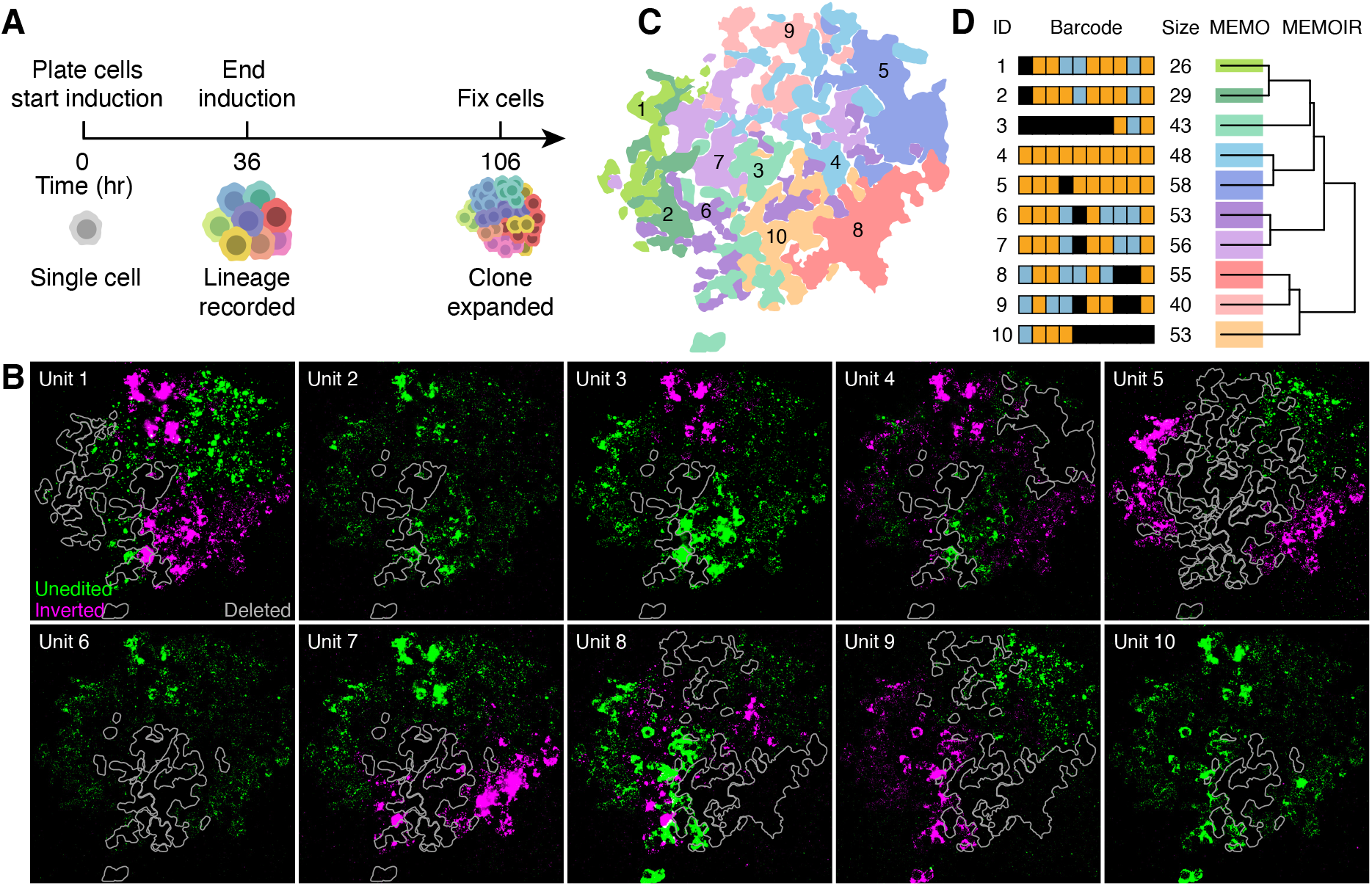
intMEMOIR enables clonal reconstruction of large colonies. (**A**) To demonstrate clonal classification and lineage reconstruction on a larger scale, we induced Bxb1 in an intMEM1 colony for 36 hours of lineage recording, followed by 70 hours of clonal expansion, fixation, and imaging. (**B**) Example of FISH readout of the intMEMOIR array in a colony. Signal for the unedited and inverted states of each unit are colored green and magenta, respectively. Clones with deleted units are outlined in gray. (**C**) Spatial distribution of the 10 clones in the colony. Cells are classified into clones based on identical edits in their intMEMOIR arrays. (**D**) Reconstruction of the early lineage of the colony. Each clone is labeled with the number of cells it contains and its corresponding color in (C). Tree reflects reconstructed lineage relationships (see Materials and Methods).

### intMEMOIR reveals spatial organization of clones and gene expression states in *D. melanogaster*

The *Drosophila melanogaster* brain provides an ideal model system to apply image-based clonal mapping *in vivo*. *Drosophila* permits rapid genetic engineering and quantitative imaging, and, while its brain development has been extensively characterized, fundamental questions about the role of lineage in fate determination remain unclear (*48*). The *Drosophila* central brain is known to develop from −100 embryonic neuroblast progenitors per hemisphere (*48*–*50*), each exhibiting a unique lineage identity, acquired largely through spatial patterning, and controlled by lineage specific transcription factors (*48*). A key step towards understanding central brain development is the capacity to uniquely label and image all distinct clones within a single organism, along with their cellular gene expression states.

To achieve such multi-clonal labeling in a single individual, we constructed a fly line that allows controllable editing and cell-type-specific readout. We cloned the intMEMOIR array downstream of a UAS promoter also expressing cyan fluorescent protein (CFP) to identify array-expressing cells, and site-specifically integrated this construct using the phiC31 system (*51*). The resulting fly line, which we term ‘*Drosophila memoiphila*’, provides a resource for general purpose lineage analysis in flies. We further incorporated a Bxb1 integrase controlled by a tightly regulated heat shock inducible promoter (*52*) to record in specific time windows. Finally, we crossed these flies to an nSyb-Gal4 driver to restrict expression of the array to neurons (Fig. 5A) (*53*).

**Fig. 5.**
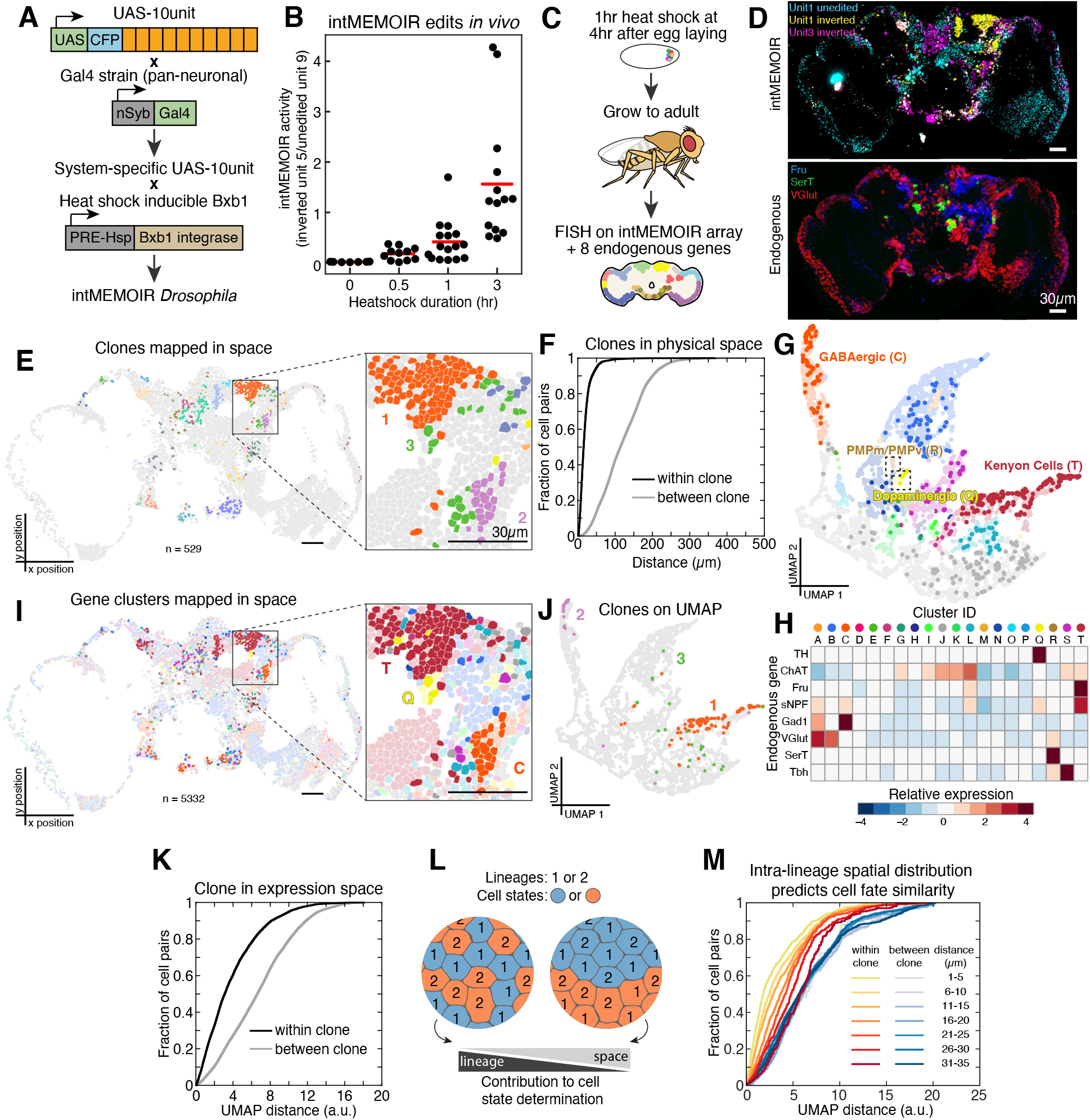
intMEMOIR connects single-cell spatial, molecular state, and lineage information in adult *Drosophila* brain. (**A**) A neuronal recording fly line was obtained by crossing *D. memoiphila* flies containing a site-specifically integrated UAS-10-unit array with an nSyb-Gal4 strain. The offspring have pan-neuronal expression of the recording array and are subsequently crossed with a tight heat shock inducible Bxb1 line to produce the final fly line. (Other Gal4 drivers could be used to analyze distinct tissues or cell types). (**B**) Editing activity, measured as the ratio of edited unit 5 over unedited unit 9, increases with heat shock duration in a dose-dependent manner (see Materials and Methods). (**C**) To achieve *in situ* analysis of cell state and clonal identity, editing was induced with a 1 hour heat shock at 4h after egg laying. This early induction aims to label neuroblasts with unique array states that can be inherited by all neuron progeny in the adult brain. Flies were then grown to adulthood, and their brains dissected and cryosectioned. Sequential rounds of FISH were used to read out the intMEMOIR array and 8 endogenous genes: tyrosine hydroxylase (TH), choline acetyltransferase (ChAT), fruitless (Fru), short neuropeptide F precursor (sNPF), glutamic acid decarboxylase (Gad1), vesicular glutamate transporter (VGlut), serotonin transporter (SerT), and tyramine β-hydroxylase (Tbh). (**D**) Example images showing single cell resolution imaging of endogenous expression and array state in the same tissue sample (scale bar, 30 μm; see Fig. S7 for additional examples). (**E**) intMEMOIR clones mapped in space. Segmented cells are colored by the 29 analyzed clones (n=529; scale bar, 30 μm). Inset highlights examples of clones that are clustered (clones 1 and 2) and dispersed (clone 3) in space. Cells outside the 29 clones are gray. Clone colors are consistent with (J). (**F**) Cells within the same clone (black) are arranged closer in physical space than cells between clones (gray). Cumulative distributions show pairwise distance between the cells. (**G**) UMAP clustering of 5,332 cells based on the expression of the 8 endogenous genes. Four clusters are annotated by inspection based on expression patterns and *in situ* localization of the majority population. Cells among the 29 analyzed clones are highlighted with saturated colors. Cluster colors are consistent between (G), (H), and (I). (**H**) Heatmap showing the relative expression, calculated as normalized Z-score, of the 8 genes in each cluster. (**I**) Gene expression clusters mapped in space. Segmented cells are colored by cluster. Cells among the 29 analyzed clones are highlighted with saturated colors (scale bar, 30 μm). The inset highlights the same cells in (E), demonstrating clones that display similar (clones 1 and 2) and mixed (clone 3) cell states. (**J**) The three represented clones mapped onto UMAP space, demonstrating examples of cells whose molecular states are correlated to their neuroblast lineage to varying degrees. (**K**) Cells within the same clone (black) are more similar in expression than cells in different clones (gray). Cumulative distributions show pairwise UMAP expression distances (Supplemental Text). (**L**) Hypothetical observations if either neuroblast lineage or spatial distribution alone dominates cell state determination in the fly brain. (**M**) Within the same clone, larger physical distances between cells correlate with greater gene expression differences (yellow to red colors). This correlation is not observed between cells of different clones (grey to blue colors). Cumulative distributions show pairwise UMAP expression distances.

To confirm intMEMOIR operation, we exposed flies to varying durations of heat shock during early development, collected adults, sectioned their brains, and read out the intMEMOIR array using sequential rounds of smFISH (*44*). Negligible levels of editing occurred without heat shock, while exposure to 37°C for 0.5h to 3h produced dose-dependent increases in editing, as quantified by analysis of two units (Fig. 5B). These results demonstrate that Bxb1 activity can be controlled in a tight, dose-dependent manner by heat shock duration (Fig. 5B).

We next sought to integrate analysis of lineage, cell state, and spatial organization in a single brain. To induce editing in neuroblasts during early embryonic development, we applied a 1 hour heat shock starting 4 hours after egg laying (Fig. 5C). We then analyzed brain sections from adults by smFISH, reading out not only the intMEMOIR array but also 8 endogenous genes selected based on their ability to identify diverse neuronal cell types (Fig. 5D and Fig. S7). Within a single brain section, we focused on the 29 clones which contained at least four cells and had at least one inverted unit (Fig. S8 and Materials and Methods). Most such clones consisted of tightly apposed cells (e.g. Fig. 5E inset, clones 1 and 2). However, a minority were more dispersed, intermingling with other clones within the section (e.g. Fig. 5E inset, clone 3) (*54*–*56*) (Fig. 5F). These results, which are consistent with previous observations (*57*, *58*), demonstrate the ability to simultaneously map the spatial arrangements of many clones in the same brain section.

We next sought to visualize the spatial distribution of gene expression states. We used UMAP (*59*) followed by density-based clustering (DBSCAN) (*60*) to reduce dimensionality and identify expression states (Fig. 5G and H). Combining gene expression with spatial location identified known cell types including GABAergic neurons, dopaminergic neurons, and Kenyon cells (Fig. 5G), and allowed us to plot their spatial distribution within the brain section (Figure 5I). Cross-referencing the spatial maps of expression states and lineage also allowed simultaneous inspection of any specific region, cell state, and lineage of interest. For example, intMEMOIR captured two out of the four known lineages of Kenyon cells in this experiment (Figure 5E and I, cluster T in the right hemisphere) (*48*). Further, by labeling distinct clones in gene expression space, we were able to visualize correlations between clonal identity and cell type similarity (Fig. 5J). This analysis revealed homogeneous clones containing a single cell type (Fig. 5J, clones 1 and 2), as well as heterogeneous clones containing multiple cell types (Fig. 5J, clone 3), consistent with previous observations (*57*). Overall, cells within the same clone tended to be more similar to one another in gene expression than cells in different clones (Fig. 5K).

In a highly structured organ such as the brain, extrinsic, spatially organized cues and intrinsic, inherited factors could both contribute to cell fate determination. To disentangle their relative contributions, we compared the similarity of gene expression states between cell pairs within the same clone to cell pairs in different clones, across a range of different spatial separations (Fig. 5M). If extrinsic cues dominate, one would expect cell state similarity to strongly correlate with spatial proximity, independent of clonal identity. By contrast, if intrinsic determinants dominate, then gene expression similarity should correlate with clonal identity, regardless of spatial separation (Fig. 5L). Here, among cell pairs drawn from distinct clones, transcriptional similarity showed little dependence on spatial distance (Fig. 5M, blue lines). However, cell pairs within the same clone showed strong cell type similarity at close distances, with a gradual relaxation of this similarity at larger distances (Fig. 5M, yellow-red lines). This result is robust to exclusion of the large, homogeneous population of Kenyon cells (cluster T, Fig. S9A), which could potentially dominate this analysis, and to other choices of gene expression distance metric (Fig. S9B). Thus, spatial proximity affects gene expression similarity within, but not between these clones, consistent with a strong role for neuroblast ancestry in cell fate determination.

## Discussion

How the lineage history of a cell impacts its future potential is a central problem in development, but challenging to systematically address in most systems. Clonal imaging methods such as MARCM revolutionized developmental analysis but are limited to a few clones per animal (*15*). Conversely, sequencing-based recording approaches such as GESTALT can provide higher throughput clonal and lineage reconstruction but do not preserve spatial information (*3*, *11*, *16*, *17*, *19*, *20*, *22*). As shown here, intMEMOIR arrays allow high-density cell-autonomous editing with image-based readout for lineage reconstruction, are compatible with smFISH gene expression measurements, and can be applied in diverse species and contexts. The genetically compact array design facilitates engineering of intMEMOIR transgenic animals, as exemplified by *D. memoiphila*, which can be crossed into any Gal4 and Bxb1 driver lines to enable recording and readout in specific cell types at desired developmental times. Here, we observed that spatial correlations of cell state in the fly central brain are strongly dependent on shared neuroblast clonal ancestry. In the future, more comprehensive analysis combining deeper lineage tree reconstructions, broader transcriptome profiling (*25*), and three-dimensional imaging should enable a finer grain view of the roles that intrinsic and extrinsic cues play in specifying the spatial arrangement of cell types. More generally, systematic incorporation of lineage information in spatially resolved cell atlases should allow researchers to explore interactions within and between clonal lineages, and to understand developmental variation between individual organisms. To achieve these goals, the memory capacity of intMEMOIR can be expanded by incorporating additional independently readable arrays. Further, because the same recording design can work with additional integrases (Fig. S2), and recent work eliminates the need for *in vivo* transcription for FISH readout (*61*), it should be possible to build intMEMOIR systems with multiple recording ‘channels’ with different integrases regulated by specific signaling pathway or transcription factor activities (*23*). intMEMOIR thus provides a foundation for understanding how each cell’s present state emerges from its spatial context and individual history.

## Acknowledgements

We thank L. Sanchez-Guardado, H. Choi, C. Calvert, G. Shin, C. Tischbirek, Y. Takei, S. Shah, and N. Pierson for technical assistance and advice. A. Askary, X. Gao, F. Horns, D. Chadley, C. Su, and other members of the Elowitz lab for critical feedback on the manuscript. A. Shur, P. Meyer, R. Lu, and J. Linton for scientific input and advice.

## Authors contributions

K.K.C., M.W.B., A.A.G., K.L.F., L.C., C.L., and M.B.E. designed research. K.K.C., M.W.B., M.C., S.Y., S.C., and N.K. performed experiments. A.A.G., M.W.B., K.K.C., T.H., C.L., and M.B.E. analyzed data, K.K.C., M.W.B., A.A.G., C.L., and M.B.E. wrote the manuscript.

## References and Notes

1. C. Blanpain, B. D. Simons, Unravelling stem cell dynamics by lineage tracing. Nat. Rev. Mol. Cell Biol. 14, 489–502 (2013).

2. K. Kretzschmar, F. M. Watt, Lineage tracing. Cell. 148, 33–45 (2012).

3. M. B. Woodworth, K. M. Girskis, C. A. Walsh, Building a lineage from single cells: genetic techniques for cell lineage tracking. Nat. Rev. Genet. 18, 230–244 (2017).

4. S.-H. S. Wu, J.-H. Lee, B.-K. Koo, Lineage Tracing: Computational Reconstruction Goes Beyond the Limit of Imaging. Mol. Cells. 42, 104–112 (2019).

5. C. S. Baron, A. van Oudenaarden, Unravelling cellular relationships during development and regeneration using genetic lineage tracing. Nat. Rev. Mol. Cell Biol. 20, 753–765 (2019).

6. D. Frumkin, A. Wasserstrom, S. Kaplan, U. Feige, E. Shapiro, Genomic variability within an organism exposes its cell lineage tree. PLoS Comput. Biol. 1, e50 (2005).

7. S. J. Salipante, M. S. Horwitz, Phylogenetic fate mapping. Proc. Natl. Acad. Sci. U. S. A. 103, 5448–5453 (2006).

8. N. Navin, J. Kendall, J. Troge, P. Andrews, L. Rodgers, J. McIndoo, K. Cook, A. Stepansky, D. Levy, D. Esposito, L. Muthuswamy, A. Krasnitz, W. R. McCombie, J. Hicks, M. Wigler, Tumour evolution inferred by single-cell sequencing. Nature. 472, 90–94 (2011).

9. S. Behjati, M. Huch, R. van Boxtel, W. Karthaus, D. C. Wedge, A. U. Tamuri, I. Martincorena, M. Petljak, L. B. Alexandrov, G. Gundem, P. S. Tarpey, S. Roerink, J. Blokker, M. Maddison, L. Mudie, B. Robinson, S. Nik-Zainal, P. Campbell, N. Goldman, M. van de Wetering, E. Cuppen, H. Clevers, M. R. Stratton, Genome sequencing of normal cells reveals developmental lineages and mutational processes. Nature. 513, 422–425 (2014).

10. M. A. Lodato, M. B. Woodworth, S. Lee, G. D. Evrony, B. K. Mehta, A. Karger, S. Lee, T. W. Chittenden, A. M. D’Gama, X. Cai, L. J. Luquette, E. Lee, P. J. Park, C. A. Walsh, Somatic mutation in single human neurons tracks developmental and transcriptional history. Science. 350, 94–98 (2015).

11. R. U. Sheth, H. H. Wang, DNA-based memory devices for recording cellular events. Nat. Rev. Genet. 19, 718–732 (2018).

12. L. S. Ludwig, C. A. Lareau, J. C. Ulirsch, E. Christian, C. Muus, L. H. Li, K. Pelka, W. Ge, Y. Oren, A. Brack, T. Law, C. Rodman, J. H. Chen, G. M. Boland, N. Hacohen, O. Rozenblatt-Rosen, M. J. Aryee, J. D. Buenrostro, A. Regev, V. G. Sankaran, Lineage Tracing in Humans Enabled by Mitochondrial Mutations and Single-Cell Genomics. Cell. 176, 1325–1339.e22 (2019).

13. H. Zong, J. S. Espinosa, H. H. Su, M. D. Muzumdar, L. Luo, Mosaic analysis with double markers in mice. Cell. 121, 479–492 (2005).

14. J. Livet, T. A. Weissman, H. Kang, R. W. Draft, J. Lu, R. A. Bennis, J. R. Sanes, J. W. Lichtman, Transgenic strategies for combinatorial expression of fluorescent proteins in the nervous system. Nature. 450, 56–62 (2007).

15. H.-H. Yu, C.-H. Chen, L. Shi, Y. Huang, T. Lee, Twin-spot MARCM to reveal the developmental origin and identity of neurons. Nat. Neurosci. 12, 947–953 (2009).

16. S. D. Perli, C. H. Cui, T. K. Lu, Continuous genetic recording with self-targeting CRISPR-Cas in human cells. Science. 353 (2016), doi: 10.1126/science.aag0511.

17. A. McKenna, G. M. Findlay, J. A. Gagnon, M. S. Horwitz, A. F. Schier, J. Shendure, Whole-organism lineage tracing by combinatorial and cumulative genome editing. Science. 353, aaf7907 (2016).

18. W. Pei, T. B. Feyerabend, J. Rossler, X. Wang, D. Postrach, K. Busch, I. Rode, K. Klapproth, N. Dietlein, C. Quedenau, W. Chen, S. Sauer, S. Wolf, T. Hofer, H.-R. Rodewald, Polylox barcoding reveals haematopoietic stem cell fates realized in vivo. Nature. 548, 456–460 (2017).

19. R. Kalhor, K. Kalhor, L. Mejia, K. Leeper, A. Graveline, P. Mali, G. M. Church, Developmental barcoding of whole mouse via homing CRISPR. Science. 361 (2018), doi: 10.1126/science.aat9804.

20. W. Tang, D. R. Liu, Rewritable multi-event analog recording in bacterial and mammalian cells. Science. 360 (2018), doi: 10.1126/science.aap8992.

21. A. Alemany, M. Florescu, C. S. Baron, J. Peterson-Maduro, A. van Oudenaarden, Whole-organism clone tracing using single-cell sequencing. Nature. 556, 108–112 (2018).

22. M. M. Chan, Z. D. Smith, S. Grosswendt, H. Kretzmer, T. M. Norman, B. Adamson, M. Jost, J. J. Quinn, D. Yang, M. G. Jones, A. Khodaverdian, N. Yosef, A. Meissner, J. S. Weissman, Molecular recording of mammalian embryogenesis. Nature. 570, 77–82 (2019).

23. K. L. Frieda, J. M. Linton, S. Hormoz, J. Choi, K.-H. K. Chow, Z. S. Singer, M. W. Budde, M. B. Elowitz, L. Cai, Synthetic recording and in situ readout of lineage information in single cells. Nature. 541, 107–111 (2017).

24. E. Lubeck, A. F. Coskun, T. Zhiyentayev, M. Ahmad, L. Cai, Single-cell in situ RNA profiling by sequential hybridization. Nat. Methods. 11 (2014), pp. 360–361.

25. C.-H. L. Eng, M. Lawson, Q. Zhu, R. Dries, N. Koulena, Y. Takei, J. Yun, C. Cronin, C. Karp, G.-C. Yuan, L. Cai, Transcriptome-scale super-resolved imaging in tissues by RNA seqFISH. Nature. 568, 235–239 (2019).

26. A. Raj, P. van den Bogaard, S. A. Rifkin, A. van Oudenaarden, S. Tyagi, Imaging individual mRNA molecules using multiple singly labeled probes. Nat. Methods. 5, 877–879 (2008).

27. I. Salvador-Martfnez, M. Grillo, M. Averof, M. J. Telford, Is it possible to reconstruct an accurate cell lineage using CRISPR recorders? Elife. 8 (2019), doi: 10.7554/eLife.40292.

28. W. M. Stark, M. R. Boocock, D. J. Sherratt, Catalysis by site-specific recombinases. Trends Genet. 8, 432–439 (1992).

29. M. C. A. Smith, R. Till, M. C. M. Smith, Switching the polarity of a bacteriophage integration system. Mol. Microbiol. 51, 1719–1728 (2004).

30. J. Bonnet, P. Subsoontorn, D. Endy, Rewritable digital data storage in live cells via engineered control of recombination directionality. Proc. Natl. Acad. Sci. U. S. A. 109, 8884–8889 (2012).

31. K. Rutherford, P. Yuan, K. Perry, R. Sharp, G. D. Van Duyne, Attachment site recognition and regulation of directionality by the serine integrases. Nucleic Acids Res. 41, 8341–8356 (2013).

32. P. C. M. Fogg, S. Colloms, S. Rosser, M. Stark, M. C. M. Smith, New applications for phage integrases. J. Mol. Biol. 426, 2703–2716 (2014).

33. Z. Xu, L. Thomas, B. Davies, R. Chalmers, M. Smith, W. Brown, Accuracy and efficiency define Bxb1 integrase as the best of fifteen candidate serine recombinases for the integration of DNA into the human genome. BMC Biotechnol. 13, 87 (2013).

34. C. A. Merrick, J. Zhao, S. J. Rosser, Serine Integrases: Advancing Synthetic Biology. ACS Synth. Biol. 7, 299–310 (2018).

35. B. P. Zambrowicz, A. Imamoto, S. Fiering, L. A. Herzenberg, W. G. Kerr, P. Soriano, Disruption of overlapping transcripts in the ROSA geo 26 gene trap strain leads to widespread expression of -galactosidase in mouse embryos and hematopoietic cells. Proceedings of the National Academy of Sciences. 94 (1997), pp. 3789–3794.

36. P. Soriano, Generalized lacZ expression with the ROSA26 Cre reporter strain. Nat. Genet. 21, 70–71 (1999).

37. M. Sadelain, E. P. Papapetrou, F. D. Bushman, Safe harbours for the integration of new DNA in the human genome. Nat. Rev. Cancer. 12, 51–58 (2011).

38. H. A. Grunwald, V. M. Gantz, G. Poplawski, X.-R. S. Xu, E. Bier, K. L. Cooper, Super-Mendelian inheritance mediated by CRISPR-Cas9 in the female mouse germline. Nature. 566, 105–109 (2019).

39. I. Espinosa-Medina, J. Garcia-Marques, C. Cepko, T. Lee, High-throughput dense reconstruction of cell lineages. Open Biol. 9, 190229 (2019).

40. P. Ghosh, A. I. Kim, G. F. Hatfull, The orientation of mycobacteriophage Bxb1 integration is solely dependent on the central dinucleotide of attP and attB. Mol. Cell. 12, 1101–1111 (2003).

41. S. D. Colloms, C. A. Merrick, F. J. Olorunniji, W. M. Stark, M. C. M. Smith, A. Osbourn, J. D. Keasling, S. J. Rosser, Rapid metabolic pathway assembly and modification using serine integrase site-specific recombination. Nucleic Acids Res. 42, e23 (2014).

42. H. Zeng, K. Horie, L. Madisen, M. N. Pavlova, G. Gragerova, A. D. Rohde, B. A. Schimpf, Y. Liang, E. Ojala, F. Kramer, P. Roth, O. Slobodskaya, I. Dolka, E. A. Southon, L. Tessarollo, K. E. Bornfeldt, A. Gragerov, G. N. Pavlakis, G. A. Gaitanaris, An inducible and reversible mouse genetic rescue system. PLoS Genet. 4, e1000069 (2008).

43. M. Iwamoto, T. Bjorklund, C. Lundberg, D. Kirik, T. J. Wandless, A general chemical method to regulate protein stability in the mammalian central nervous system. Chem. Biol. 17, 981–988 (2010).

44. H. M. T. Choi, M. Schwarzkopf, M. E. Fornace, A. Acharya, G. Artavanis, J. Stegmaier, A. Cunha, N. A. Pierce, Third-generation in situ hybridization chain reaction: multiplexed, quantitative, sensitive, versatile, robust. Development. 145 (2018), doi: 10.1242/dev.165753.

45. D. F. Robinson, L. R. Foulds, Comparison of phylogenetic trees. Math. Biosci. 53, 131–147 (1981).

46. K. Sugino, J. Garcia-Marques, I. Espinosa-Medina, T. Lee, Theoretical modeling on CRISPR-coded cell lineages: efficient encoding and optimal reconstruction. bioRxiv (2019), p. 538488.

47. J. M. Kebschull, A. M. Zador, Cellular barcoding: lineage tracing, screening and beyond. Nat. Methods. 15, 871–879 (2018).

48. T. Lee, Wiring the Drosophila Brain with Individually Tailored Neural Lineages. Curr. Biol. 27, R77–R82 (2017).

49. R. Urbach, G. M. Technau, Neuroblast formation and patterning during early brain development in Drosophila. Bioessays. 26, 739–751 (2004).

50. S. R. Spindler, V. Hartenstein, The Drosophila neural lineages: a model system to study brain development and circuitry. Dev. Genes Evol. 220, 1–10 (2010).

51. A. C. Groth, M. Fish, R. Nusse, M. P. Calos, Construction of transgenic Drosophila by using the site-specific integrase from phage phiC31. Genetics. 166, 1775–1782 (2004).

52. A. Akmammedov, M. Geigges, R. Paro, Single vector non-leaky gene expression system for Drosophila melanogaster. Sci. Rep. 7, 6899 (2017).

53. O. Riabinina, D. Luginbuhl, E. Marr, S. Liu, M. N. Wu, L. Luo, C. J. Potter, Nat. Methods, in press.

54. K. Dumstrei, F. Wang, C. Nassif, V. Hartenstein, Early development of the Drosophila brain: V. Pattern of postembryonic neuronal lineages expressing DE-cadherin. J. Comp. Neurol. 455, 451–462 (2003).

55. K. Ito, W. Awano, K. Suzuki, Y. Hiromi, D. Yamamoto, The Drosophila mushroom body is a quadruple structure of clonal units each of which contains a virtually identical set of neurones and glial cells. Development. 124, 761–771 (1997).

56. S.-L. Lai, T. Awasaki, K. Ito, T. Lee, Clonal analysis of Drosophila antennal lobe neurons: diverse neuronal architectures in the lateral neuroblast lineage. Development. 135, 2883–2893 (2008).

57. M. Ito, N. Masuda, K. Shinomiya, K. Endo, K. Ito, Systematic analysis of neural projections reveals clonal composition of the Drosophila brain. Curr. Biol. 23, 644–655 (2013).

58. H.-H. Yu, T. Awasaki, M. D. Schroeder, F. Long, J. S. Yang, Y. He, P. Ding, J.-C. Kao, G. Y.-Y. Wu, H. Peng, G. Myers, T. Lee, Clonal development and organization of the adult Drosophila central brain. Curr. Biol. 23, 633–643 (2013).

59. E. Becht, L. McInnes, J. Healy, C.-A. Dutertre, I. W. H. Kwok, L. G. Ng, F. Ginhoux, E. W. Newell, Dimensionality reduction for visualizing single-cell data using UMAP. Nat. Biotechnol. (2018), doi: 10.1038/nbt.4314.

60. M. Ester, H.-P. Kriegel, J. Sander, X. Xu, Others, in Kdd (1996), vol. 96, pp. 226–231.

61. A. Askary, L. Sanchez-Guardado, J. M. Linton, D. M. Chadly, M. W. Budde, L. Cai, C. Lois, M. B. Elowitz, In situ readout of DNA barcodes and single base edits facilitated by in vitro transcription. Nat. Biotechnol. 38, 66–75 (2020).

62. S. Yamaguchi, Y. Kazuki, Y. Nakayama, E. Nanba, M. Oshimura, T. Ohbayashi, A method for producing transgenic cells using a multi-integrase system on a human artificial chromosome vector. PLoS One. 6, e17267 (2011).

63. S. Berg, D. Kutra, T. Kroeger, C. N. Straehle, B. X. Kausler, C. Haubold, M. Schiegg, J. Ales, T. Beier, M. Rudy, K. Eren, J. I. Cervantes, B. Xu, F. Beuttenmueller, A. Wolny, C. Zhang, U. Koethe, F. A. Hamprecht, A. Kreshuk, ilastik: interactive machine learning for (bio)image analysis. Nat. Methods. 16, 1226–1232 (2019).

64. I. T. Jolliffe, J. Cadima, Principal component analysis: a review and recent developments. Philos. Trans. A Math. Phys. Eng. Sci. 374, 20150202 (2016).

