## Supplementary Materials for "Imaging cell lineage with a synthetic digital recording system"

**This PDF file includes:**

Materials and Methods
Supplementary Text
Figs. S1 to S9
References

**Other Supplementary Materials for this manuscript include the following:**

Tables S1 and S2 (.xlsx)

### Materials and Methods

Plasmids preparation

Constructs were cloned using standard methods. Due to the repetitive sequence, inverted *attP*s were difficult to amplify *in vitro*, therefore PCR-based cloning methods were avoided for these regions. Mammalian constructs involving serine integrases Bxb1, phiC31, R4, and TP901 were cloned from, or used directly as, plasmid gifts from Mitsuo Oshimura (*62*). All constructs reported in this manuscript are listed in Table S1, and sequence maps for constructs generated for the intMEMOIR system are available at https://data.caltech.edu/.

Tissue culture

All tissue culture experiments were done with E14 mouse embryonic stem (mES) cell line (ATCC catalog number CRL-1821). Cells were cultured in humidified chambers at 37ºC and 5% $CO_2$, with filtered media composed of GMEM (Sigma), 15% FBS, PSG (100 units/mL penicillin, 100 µM/mL streptomycin, 2 mM L-glutamine) (ThermoFisher), 1mM sodium pyruvate (ThermoFisher), 1X Minimum Essential Medium Non-Essential Amino Acids (MEM NEAA, ThermoFisher), and 100µM 2-Mercaptoethanol (ThermoFisher), with 1,000 units/mL Leukemia Inhibitory Factor (LIF, Millipore) added after filtering. Cells were maintained on polystyrene plates coated with 0.1% gelatin.

Transient transfection

Transient transfections were performed with Lipofectamine LTX and PLUS reagent (ThermoFisher) overnight. To reduce the presence of untransfected cells in downstream analysis, constructs are either cotransfected with fluorescent markers (mTagBFP) or antibiotic resistance plasmids (puromycin), the latter followed by two days of selection with 1µg/mL puromycin.

Flow cytometry

Flow cytometry was performed two days after transfection on CytoFlex (Beckman Coulter). Cells were lifted from the plate with StemPro Accutase (ThermoFisher) and resuspended in buffer made of HBSS, 2.5mg/mL BSA, and 1mM EDTA. They were then filtered through a 40 µm cell strainer prior to flow cytometry. These experiments, including their respective transfections, were conducted in triplicate (Fig. 2C to E).

Flow cytometry data were analyzed using the EasyFlow Matlab program developed by Yaron Antebi, the latest version available at https://antebilab.github.io/easyflow/. We gated for single cells using forward and side scatter (FSC and SSC), and gated for cells expressing high levels of a cotransfection marker. More specifically, we selected cells with mTagBFP fluorescence > 15,000 (a.u.) to enrich for the transfected population in downstream analysis. We then determined the median Citrine and mCherry fluorescence, averaged over the experimental triplicates, and background subtracted the fluorescence detected in the no integrase negative control. The resulting values were normalized and plotted for comparison, with the error bars representing normalized standard error of the mean (SEM).

intMEM1 cell line construction

To construct intMEM1, we began by integrating a landing pad containing FRT sites into the TIGRE locus using Cas9-mediated homologous recombination (construct modified from (*42*)), followed by selection with 10 µg/mL blasticidin. After selecting a clone with correct integration, we introduced constitutive pEF1α-Tet3G through PiggyBac transposition (System Biosciences) followed by 1 µg/mL puromycin treatment, and again selected for a single clone. We then inserted TRE-Bxb1-ecDHFR into the TIGRE landing pad with FlpE recombinase and 100 µg/mL hygromycin selection. The resulting polyclonal line was transfected with the 10-unit intMEMOIR array targeted to the Rosa26 locus through Cas9-mediated homologous recombination, 500 µg/mL geneticin selection, and monoclonal selection. Finally, we increased the fluorescence of these cells for time-lapse movie tracking by integrating PGK-mTurquoise2 by PiggyBac, followed by blasticidin and a second round of hygromycin selection. The final round of monoclonal selection resulted in the intMEM1 cell line. All monoclonal selections involving site-specific integrations were screened with PCR.

Time-lapse imaging for ground truth lineage

Cells were plated on glass bottom 24-well plates (Eppendorf) coated with 20 µg/mL of Laminin-511 (BioLamina) overnight. Approximately 6,000 intMEM1 cells were seeded onto the coated wells, along with 18,000 parental E14 cells to increase cell density to support growth and survival. Media was changed prior to the start of the movie to remove any unattached cells. Under a microscope equipped with an environmental chamber, intMEM1 cells were selected for time-lapse imaging by their mTurquoise2 fluorescence. Imaging then began with the induction of 10 µM TMP (to block the DHFR degron) and 100 ng/mL doxycycline (to activate the TRE3G promoter). Inducers were omitted in negative control samples. For each position, images were acquired every 15 minutes in both the visible light (DIC) and fluorescent (CFP) channels. 36 hours after the start of the movie we halted induction by washing off the induction media and replacing it with regular culture media. 54 hours after the start, we terminated time-lapse imaging and promptly fixed the sample at room temperature with 4% formaldehyde in PBS for 5 minutes, followed by smFISH protocol (below).

Constructing ground truth lineage

Ground truth lineage trees were constructed by manually tracking the cells in the time-lapse images using a modified version of the EasyTrack software developed by Yaron Antebi. Cells were primarily followed by their CFP fluorescence. Ground truth trees could begin at either the one or two cell stage depending on the colony's cell cycle at the start of image acquisition, and were only rooted at the two cell stage if the parent cells in question were likely sisters based on proximity, cell morphology, CFP intensity, as well as their cell movements and cycles in the subsequent frames. Ground truth trees for all colonies were outputted as Newick strings (Table S2).

Hybridization Chain Reaction (HCR) smFISH

For HCR smFISH in tissue culture, fixed cells were washed with PBS after formaldehyde fixation, followed by permeabilization in 70% RNase-free ethanol for durations ranging from 4 hours (4$^{o}$C) to overnight (-20$^{o}$C). Permeabilized cells were washed with 20% formamide wash buffer in 2X SSCT at room temperature for 5 minutes and pre-hybridized in 30% probe hybridization buffer at 37$^{o}$C for 30 minutes. Primary probe hybridizations and hairpin amplifications were then carried out as previously described for HCR v3.0 (Molecular Instruments) (*44*). Cells were imaged in 5X SSCT.

Between hybridization rounds, probes were removed via DNase I treatment (Roche). Briefly, cells were washed with 1X DNase buffer, followed by incubation with 1 Kunitz unit/µL DNase I in 1X buffer for 2 to 4 hours at 37$^{o}$C. Digestion was ended by washing the cells 3 times with 30% probe wash buffer, incubating the final wash for 15 minutes at 37$^{o}$C. Finally, cells were washed once with 5X SSCT before the pre-hybridization step for the next round of smFISH.

Antibody staining

Upon completion of all rounds of smFISH readout and a final round of DNase I treatment to remove any smFISH signals, cells underwent antibody staining for membrane markers E-cadherin and β-catenin to facilitate segmentation. Immunostaining was performed following standard protocols, with most incubation and washing steps carried out on a gentle rocker. Briefly, samples were blocked with blocking buffer made in PBS (5% BSA, 1% DMSO, and 0.2% Triton X-100) for 1 hour at room temperature. They were then incubated with primary antibodies E-cadherin (R&D Systems, AF648, 1:20) and β-catenin (Abcam ab6301clone15B8, 1:750) overnight at 4$^{o}$C. The following day, they were washed 5 times with PBST for 5 minutes each, then incubated with secondary antibodies (donkey anti-goat IgG 647 A21447, and donkey anti-mouse IgG 488 A21202, respectively) diluted 1:1000 in blocking buffer for 3 hours at room temperature. Finally, samples were washed 5 times with PBS for 5 minutes each, followed by imaging in fresh PBS.

Analysis of smFISH readout in mES cells

To segment individual cells and identify their array edit states, we used a custom analysis pipeline in Matlab. For cell segmentation, immunofluorescence images of the membrane were first preprocessed with Ilastik (*63*) to generate a membrane probability map. The positions of cells in the final frame of the ground truth lineage analysis were used as watershed seeds overlaid on the membrane probability maps. The resulting segmented images were visually examined and manually curated.

For array state determination, Ilastik was first used for pixel classification of the fluorescence signal of the individual HCR smFISH images, and then the centers of the mRNA dots were determined using a Laplacian of Gaussian filter in Matlab. Barcode mRNA locations were called when multiple units localized to the same spot. Each barcode mRNA state was determined by looking at the binary state of each of the twenty unit smFISH images. All of the barcodes located in each cell were used to generate a consensus barcode state. Cells with fewer than 50 detected units were excluded from analysis.

Calculation of mutual information between recording units

We pooled the empirical data from all 1,453 cells and built a frequency matrix $\Gamma_{3\times 10}$ representing each of the 10 recording units and the observed frequency of each one of the three possible intMEMOIR states which define the distribution $P(x)$ per site. For each pair of sites, we then computed the joint distribution $P(x, y)$ from the observed frequencies of pairs of states e.g. $[(1,1),(1,0),(1,2),]$. We then combined the probabilities in $\Gamma$ with the joint distribution to build a matrix of pairwise Mutual Information using Shannon's formula, using $log_3$ to normalize the maximum entropy of a single unit to 1 trit.

Lineage analysis of large mES colony

Cells were plated on glass bottom 24-well plates (Eppendorf) coated with 20 µg/mL of Laminin-511 overnight. Approximately 1,000 intMEM1 cells were seeded onto the coated wells and induced with 10 µM TMP and 100 ng/mL doxycycline, along with 9,000 parental E14 cells to increase cell density to support growth and survival. Induction lasted 36 hours, followed by approximately 70 hours of growth with no induction. Media was changed daily, and the cells were fixed at the end of the experiment with 4% formaldehyde in PBS for 5 minutes, followed by smFISH protocol described above. Clone boundaries and barcode analysis for this colony were analyzed by hand.

*D. memoiphila* fly line generation

Fly lines containing UAS-Ceru-10unit and PRExpress-Bxb1-hsp70pA were site-specifically integrated into the attp2 and VK27 sites, respectively, using phiC31 (Bestgene Inc.). Flies with the 10-unit array were first crossed with an nSyb-Gal4 line (R57C10-Gal4, attp40, Bloomington Drosophila stock center) for pan-neuronal expression of the intMEMOIR array. The offspring were then crossed with the PRExpress-Bxb1-hsp70pA to generate the line capable of autonomous recording for downstream experiments.

D. memoiphila characterization

To determine if we could tune the edits in *D. memoiphila* embryo and read out the results in adult fly brains, we placed parents from the PRExpress-Bxb1-hsp70pA x nSyb-Gal4 x UAS-10unit/TM3 cross in fresh vials overnight at 25ºC to collect eggs. The embryos were heat shocked in 37ºC water bath approximately 3-4 hours later for 30 minutes, 1 hour, and 3 hours for the respective samples. Negative control samples were always kept at 25ºC and not heat shocked. The resulting adult flies were sacrificed, and their brains were dissected in PBS and fixed in 4% paraformaldehyde (PFA) in PBS for 20 minutes. Samples were then washed 3 times with PBS for 10 minutes, transferred to Optimal Cutting Temperature (OCT) compound, and frozen on dry ice. Samples were cut into 20 µm-thick sections on a cryostat and transferred onto silane-coated slides, followed by post-fix with 4% PFA in PBS for 15 to 25 minutes, 3 rinses with PBS, 1 rinse with 70% ethanol, and permeabilized in 70% ethanol overnight at 4ºC. Tissues were then cleared with 8% SDS for 5 minutes at room temperature, rinsed once with PBS, then 3 times with 70% ethanol. intMEMOIR array states were then read out with HCR v3.0 as described (Molecular Instruments) (*44*).

To eliminate error or bias that may result from image alignment between hybridizations, we evaluated intMEMOIR activity by calculating the ratio of inverted unit 5 to unedited unit 9: two efficiently edited units (as indicated by mES data, Fig. 2I) that were probed in the same round of hybridization. Each data point in Fig. 5B corresponds to one imaging position.

intMEMOIR labeling of early neuroblast lineages

To label neuroblast lineages at early embryonic stages, parents from the PRExpress-Bxb1-hsp70pA x nSyb-Gal4 x Bxb1 UAS-10unit/TM3 cross were placed in fresh vials for 1 hour at 25ºC to collect freshly laid eggs. The parents were removed, and 4 hours later the embryos were heat shocked at 37ºC for 1 hour. The resulting adult flies, up to 1 week old, were incubated at 29ºC overnight to enhance activity of the Gal4 transcription factor on the UAS promoter prior to brain collection and cryosection, followed by smFISH readout (below).

smFISH readout in *D. melanogaster* brain section

To simultaneously examine spatial organization, cell state, and lineage information in the same tissue (Fig. 5D to M), the dissected fly brains were cryosectioned and attached onto coverslips treated with 1% bind-silane and poly-D-lysine, and the resulting samples prepared and analyzed with sequential, automated rounds of smFISH in a manner similar to previously described (*25*). Briefly, sections were post-fixed with 4% PFA at room temperature for 15 minutes, followed by three PBS washes. They were then permeabilized in 70% ethanol (either at 4˚C overnight or at room temperature for 2 hours), cleared with 8% SDS in 1X PBS for 20 minutes at room temperature, then washed with 70% ethanol prior to primary probe hybridization overnight at 37˚C. After hybridization, samples were washed with 2X SSC for 3 times, incubated in 40%

formamide in 2X SSC for 30 minutes at 37°C, followed by 3 additional rounds of 2X SSC wash. They were then stained with 100 μg/mL Concanavalin A-488 (ThermoFisher) in PBS with 0.1% BSA and 0.1% Triton X-100 for more than 5 hours at room temperature to facilitate segmentation in downstream analysis. After staining, the sample was washed three times with PBS plus 0.5% Triton X-100 (with an extended 5 minute incubation for the final wash), and stained with 10 μg/ml DAPI in 4X SSC for 15 seconds. Next, an anti-bleaching buffer solution made of 10% (w/v) glucose, 1:100 diluted catalase, 0.5 mg/ml glucose oxidase and 50 mM pH 8 Tris-HCL in 4X SSC was flowed through the samples. Finally, we proceeded with sequential rounds of FISH to read out the intMEMOIR array and 8 endogenous genes using the automated imaging and fluidics delivery system described in (*25*).

Analysis of smFISH readout in *D. melanogaster* brain section

Fly cells were segmented manually. A custom Matlab program was used to determine the barcode state, as with the mES cells, except that the Laplacian of Gaussian filter was performed directly on the smFISH images without preprocessing. Clones which had at least one barcode inversion and at least 4 cells were chosen for downstream analysis. The average fluorescence value of pixels in the segmented cell was used as the endogenous gene expression value.

Gene expression analysis in brain section

The brain data set comprises gene expression, location, and intMEMOIR state for 5,332 individual cells. For each cell, we recorded the expression of 8 endogenous genes. To investigate the structure of the gene expression space, we constructed a gene expression matrix $M_{m \times n}$, where $n$ = 8 genes, and $m$ = 5,332. Based on this matrix, the analysis pipeline we built to delineate gene expression clusters consists of several steps: 1. Scaling gene expression using a z-transform, such that all genes have mean=0 and standard deviation=1. 2. Denoising data by applying PCA (*64*) to the scaled data and retaining principal components accounting for 80% of the total variance (6 PCs). 3. The 6-dimensional data were then clustered using DBSCAN (*60*) (sklearn DBSCAN with eps = 0.3) which resulted in a total of 20 distinct clusters. 4. For visualization, the 6-dimensional data set was transformed and projected into 2 dimensions using UMAP (*59*) (default parameters, Python UMAP v0.3.10). 5. Finally, we mapped the gene expression clusters (as color labels) into either the UMAP space (Fig. 5G) or physical space (Fig. 5I).

Determining the relationship between clonality, physical distance, and gene expression distance

The physical distance was calculated as the Euclidean distance between all pairs of barcoded cells chosen for analysis. This data set was then divided into two groups: pairs within the same clone or pairs from two different clones, and plotted as a cumulative histogram (Fig. 5F, 'within clone' and 'between clone', respectively). For gene expression space, the Euclidean distance was calculated between cell pairs using the UMAP coordinates (Fig. 5K and Fig. S9A). To disentangle the relative contribution of physical distance and lineage to gene expression, the aforementioned data was further binned by the physical Euclidean distance between cell pairs (Fig. 5M). Pearson correlation was also used as an alternative metric for gene expression distance (Fig. S9B).

### Supplementary Text

Here we describe procedures for two types of lineage analysis (Fig. 1A). First, we discuss assignment of individual cells to clones, i.e. groups of cells that share a common ancestor at the time of editing (clonal classification). Second, we discuss the hierarchical assignment of cells or clones into multi-generational lineage trees (lineage tree reconstruction). In both cases, we describe an analytical framework and experimental validation using the data in Figure 3. These data were obtained from experiments in which Bxb1 was expressed for ~3 generations, followed by an additional ~1-2 generations of clonal expansion without Bxb1 induction (Fig. 2G).

Clonal classification

intMEMOIR can classify cells into clones based on shared array state inherited from a common ancestor that was uniquely labeled at a specific point in the past. Clonal analysis can be used both to address specific biological questions, and to provide the ‘leaves’ of more detailed lineage tree reconstruction (below).

Ideally, clonal classification should group cells in such a way that each cell is more closely related to other cells in its own group than to any cell in other groups. In general, a given lineage tree can generate multiple, distinct clonal classifications depending on which edits occurred at what point in the tree. For example, the tree in Figure S3 could show multiple distinct sets of unique edit patterns (colors), all consistent with the true lineage.

An experimental clonal classification is made in a straightforward way by grouping cells with identical edit patterns into putative clones. To assess the accuracy of such a clonal classification, we must first determine if it is consistent with the ground truth lineage tree observed by direct time-lapse imaging (Fig. 3A), and, second, quantify the number and types of classification errors, if any (Fig. 3C).

The following algorithm assigns an accuracy score to a given putative clone, labeled R. To do so, it considers all subtrees (partitions) of the ground truth lineage tree and asks whether any subtree exactly matches the inferred clone. If such a subtree exists, then the clonal classification is considered accurate. If not, we identify the subtree that most closely matches the clone, which we label as S, and quantify its deviation from the putative clone. This deviation is computed by first classifying each cell in S as either a true positive (appears in both S and R), a false positive (appears in R but not S), or a false negative (appears in S, but not R). We then count the number of cells in each of these three categories and compute a clone score:

$$\text{score} = \frac{TP}{TP + FP + FN}$$

Here, $TP$, $FP$, and $FN$ denote the number of cells that are true positive, false positive, or false negative, respectively. A higher score indicates a higher fraction of true positive cells (greater accuracy). Results from this analysis are plotted in Fig. 3D.

Lineage reconstruction

To reconstruct a multi-generation lineage tree from observed edit patterns, we first develop a relatedness metric for pairs of cells (or clones) based on their edit patterns. The metric is based on the likelihood of a sister relationship. We then use this metric to reconstruct lineage trees in such a way that cells that score higher on this sister likelihood metric are grouped more closely together on the reconstructed tree. Finally, we validate this procedure and quantify its accuracy.

To develop the metric, we start by modeling the molecular events that generate the final edit patterns. We assume each unedited memory element can be edited stochastically at a constant, empirically determined rate per cell generation per memory element, denoted $\mu_k$, where $k = 1..10$ indexes the memory element within the array (see Fig. 2I). We also incorporate the empirical transition probabilities for each of the two possible outcomes of each state (Fig. 2I). We then define the probability distribution $P_g$ for observing each of the 10-unit array states that occur in the colony of interest starting from an unedited array, after $g$ generations. $P_g$ is computed for each cell in the colony, and represents the probability of observing that cell's specific array state, independent of the states of other cells in the colony.

Next, we define the pairwise distance metric for a single memory element. We denote the conditional probability of observing any two specific memory states in a pair of sister cells as $P_g^{sis}(i,j)$, where $i$ and $j$ index two specific, different cells $(i \neq j)$. To convert this probability into a distance metric, we need to normalize it by comparing the likelihood of observing these two memory states in a pair of sister cells to the likelihood of them occurring independently in two unrelated cells. That is, we define the distance metric as

$$d_{i,j} \equiv \frac{P_g(i)P_g(j)}{P_g^{sis}(i,j)}.$$

Memory units edit independently (Fig. 2J). Therefore, it is possible to extend this distance metric for a single memory element to the level of a complete array in a straightforward manner, by replacing the single unit probabilities with products over all the units:

From this we obtain the $K = 10$ unit array distance metric:

$$d_{i,j}^{array} \equiv \frac{\prod_{k=1}^{10} P_g(i)_k P_g(j)_k}{\prod_{k=1}^{10} P_g^{sis}(i,j)_k}$$

*Deriving probability distributions for the intMEMOIR system*

This distance metric is independent of many details of the recording system. To apply it to intMEMOIR data, we first need to derive expressions for the distributions $P_g$ and $P_g^{sis}$. The recording units have an initial state, denoted *1*, that can be edited irreversibly into either of two states, denoted *0* and *2*. The probability that a given unit is edited during a cell division is $\mu_k$, and the probability that no edit happens during a cell division is $(1 - \mu_k)$. For simplicity, we first derive $P_g$ assuming only two possible states: unedited $(1)$ and edited $(0)$. For a given unit, the probability that no editing happens for *g* generations (cell divisions) is then

$$P_g(1) = (1 - \mu)^g \quad (1)$$

The probability that an edit occurred at some point in the past is defined by the geometric distribution:

$$P_g(0) = \sum_{g=1}^{G} (1 - \mu)^{g-1} \mu \quad (2)$$

This expression considers all possible times at which the edit could have happened, e.g. in the first generation, the second generation, or even in the last generation. Once the edit occurs, the unit can no longer be edited. (Below, we will extend this analysis to the case of multiple edit outcomes.)

By applying the geometric series, we can show that $P_g$ is well defined as a probability distribution for all values of *g* such that:

$$P_g(0) + P_g(1) = 1 \quad (3)$$

We derive Eq. 3 by first expanding Eq. 2:

$$P_g(0) = \mu + (1 - \mu)\mu + (1 - \mu)^2\mu + (1 - \mu)^3\mu + ... + (1 - \mu)^{g-1}\mu$$

Combining eqs. 1 and 2 we obtain the total probability:

$$P_g(0) + P_g(1) = \mu[1 + (1 - \mu) + (1 - \mu)^2 + ... + (1 - \mu)^{g-1}] + (1 - \mu)^g$$

We then use the following identity for the geometric series:

$$\sum_{k=0}^{n-1} qw^k = q\frac{1 - w^k}{1 - w}$$

By setting $q = 1$, $w = 1 - \mu$ and $k = g$, we obtain:

$$P_g(0) + P_g(1) = \mu \frac{1 - (1-\mu)^g}{\mu} + (1-\mu)^g = 1$$

Which shows that $P_g$ is well-defined for all values of $g$.

*Three-state model*

We now extend the model by considering three possible edit outcomes: $\{1,0,2\}$. The probabilities of observing the recording unit in each of three possible states at generation $g$ become:

$$P_g(1) = (1-\mu)^g$$

$$P_g(0) = \sum_{g=1}^{G} (1-\mu)^g \mu\alpha$$

$$P_g(2) = \sum_{g=1}^{G} (1-\mu)^g \mu(1-\alpha) \tag{4}$$

Here, $\alpha$ denotes the probability of an edited unit going to state *0* and $1-\alpha$ is the probability of it reaching state $2$. Note that, in a similar way, this framework could also be generalized to larger numbers of editing outcomes. The transition probability distribution $P(z_{g-1} \to i)$ represents all the ways in which an individual unit can change state during a single cell division cycle:

$$P(1 \to 1) = 1 - \mu$$

$$P(1 \to 0) = \mu\alpha$$

$$P(1 \to 2) = \mu(1-\alpha) \tag{5}$$

$$P(0 \to 0) = 1$$

$$P(2 \to 2) = 1$$

The probabilities of all other transitions are zero, consistent with the irreversibility of intMEMOIR editing. Further, these transition rates are assumed to be time independent.

*Sister likelihood*

We now have all elements necessary to compute the sister likelihood scores that are the basis of our distance metric. We calculate the conditional probability that an unobserved parental state $z$ in the previous generation transitioned into the observed states $i, j$. For simplicity, first consider just a single cell state, $i$:

$$P_g(i|z_{g-1}) = P(z_{g-1} \to i|z_{g-1}) P_{g-1}(z) \tag{6}$$

Where $P_{g-1}(z)$ is the probability of observing the parental state $z$ in the previous generation and can be calculated using Eq. 4. Considering now two cells $i, j$, this transition probability becomes the joint distribution:

$$P_g(i,j|z_{g-1}) = P(z_{g-1} \to i|z_{g-1})P(z_{g-1} \to j|z_{g-1})P_{g-1}(z) \tag{7}$$

Eq. 7 provides the probability that the observed states $i, j$ came from the unobserved parental state $z$. Since we don't actually observe $z$, we need to account for all possible parental states to obtain the total sister probability $P_g^{sis}$:

$$P_g^{sis}(i,j) = \sum_{z=0}^{2} P_g(i,j|z_{g-1}) \tag{8}$$

Finally, considering an array of $k$ independent units we extend this and calculate the product:

$$P_g^{sis}(C_i, C_j) = \prod_{k=1}^{10} P_g^{sis}(i,j)_k \tag{9}$$

Where $C_i, C_j$ represent a specific pair of array states, e.g. in two different cells or clones.

*Hierarchical lineage tree*

The calculations above enable us to compute the pairwise distance matrix $d_{i,j}^{array}$, defined above, for any actual data set. As the final step in reconstruction, we built a dendrogram from $d_{i,j}^{array}$ by applying divisive clustering, a top-down approach in which all observations start in one cluster, and two-way splits are performed recursively as one moves down the lineage tree, terminating at the leaves (individual cells or clones). Divisive clustering was implemented with the *DIANA* function from the **R** *cluster* package. Examples of the resulting trees are shown in Figures 3D to 3F. A complete list of reconstructed trees is provided in Table S2 in the Newick tree format.

*Assessing tree accuracy*

To quantitatively assess reconstruction accuracy compared to ground truth (Fig. 3H), we used the Robinson-Foulds distance metric, as implemented in the **R** *phytools* package, to compute the difference between the reconstructed and ground truth trees. For this analysis, all cells sharing the same array state are collapsed into a single clonal tree "leaf." Occasionally, analysis of ground truth trees revealed convergent edits producing identical array states in distantly related cells (false positive events; see discussion of clonal analysis, above). These events prevent one from unambiguously collapsing identical array states in the ground truth tree. In these cases, we randomly retain one of the array states.

Calculation of entropy in colonies

In applications where no ground truth is available, we would like to develop a predictive metric that could be used to enrich for colonies that are likely to reconstruct with greater accuracy. Multiple variables, including spatial arrangements of cells or morphological similarity could in principle be informative. However, we reasoned that the most useful and generalizable metric would be one based only on the observed edit patterns, since this information should be available in all applications independent of systems-specific biological features.

Shannon's entropy provides an ideal metric to quantify information content in a discrete data set such as a list of edit states. To apply it to colonies, we pooled the edit states from all cells within each colony. We then constructed a $10 \times 3$ matrix, $\Gamma$ representing the frequency of observing each of the 10 recording units in each of the 3 edit states. We can then apply Shannon's formula to each unit to obtain it's individual entropy, e.g. $H_k = -\sum_{i=0}^{2} \Gamma_{i,k} \log \Gamma_{i,k}$ is the entropy of the $k^{th}$ unit. The total entropy of the colony is then obtained by adding the entropies of the individual sites: $H_{colony} = \sum_{k=1}^{10} H_k$. Finally, we scaled the entropy by the fraction of edited sites in the colony, $\pi$, to obtain an informative score of expected reconstruction quality. We confirmed that selecting colonies with higher normalized entropy enriched for better reconstruction (Fig. 3H and Fig. S5).

### Supplementary Figures

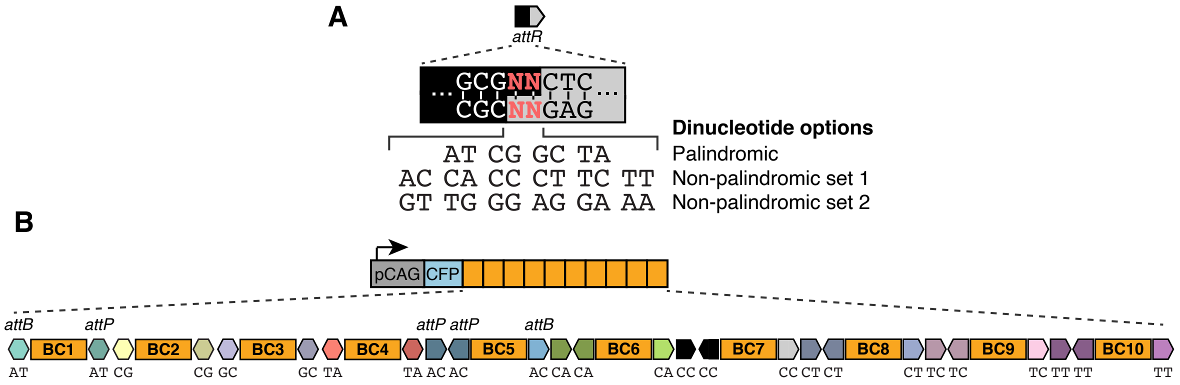


**Fig. S1. 10 central dinucleotide variants in the *att* site enable the arrangement of 10 independent memory units in one array.**

(**A**) The central dinucleotide (red) of *attP/B* form base pairs during recombination, dictating specificity and orthogonality of the sites. In principle, out of the 16 possible dinucleotide combinations, 10 could confer orthogonality in an array: four are palindromic, and the two sets of six non-palindromic dinucleotides are reverse complements of one another, so only one set could be used. (**B**) Schematic of the 10-unit array as designed with annotated *att* sites and barcodes (BC).

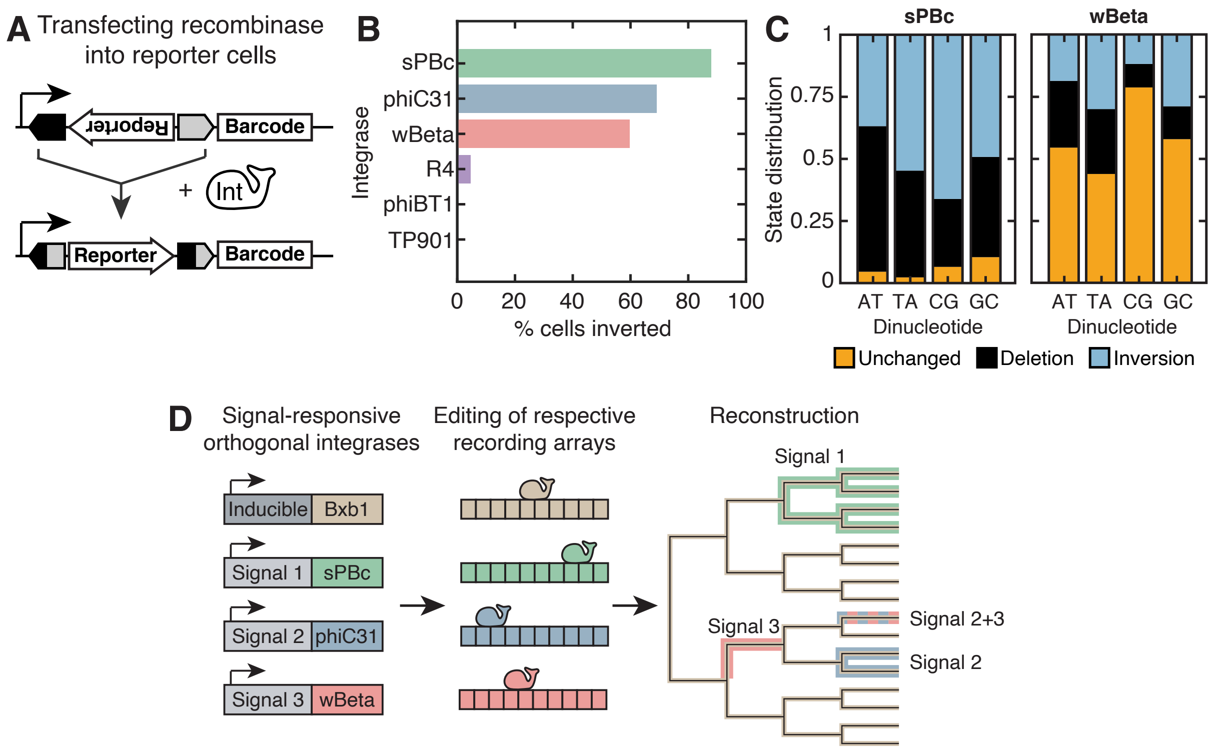


**Fig. S2. Additional members of the serine integrase family function in mES cells.**
(**A**) To assess the activity of different serine integrases in mES cells (*33*), we integrated reporter constructs with an *attP/B* flanked unit followed by a barcode. Active integrase inverts the unit upon transfection, which can be detected via smFISH. (**B**) Percentage of cells with inverted reporter after transfection. sPBc, phiC31, and wBeta are active in mES cells; R4 show weak activity, and no activity was detected for phiBT1 and TP901. (**C**) Additional serine integrases can mediate inversion and deletion between *att* sites with palindromic dinucleotides. sPBc and wBeta reporter cell lines with a 4 unit array were transfected with their corresponding integrases, and their relative edit frequencies analyzed via smFISH. (**D**) Schematic illustrating simultaneous lineage tracking and signal recording using orthogonal serine integrases.

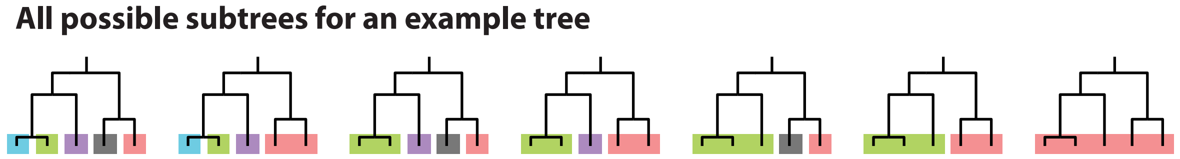


**Fig. S3. A given lineage tree can generate multiple, distinct clonal classifications.** Seven possible subtree assignments can accurately describe this simple five-cell tree, where all cells within a clone are more closely related to each other than to any cell outside of the clone. In one extreme, each individual cell can be classified as its own clone. At the other extreme, in the absence of editing, all cells are grouped into a single clone.

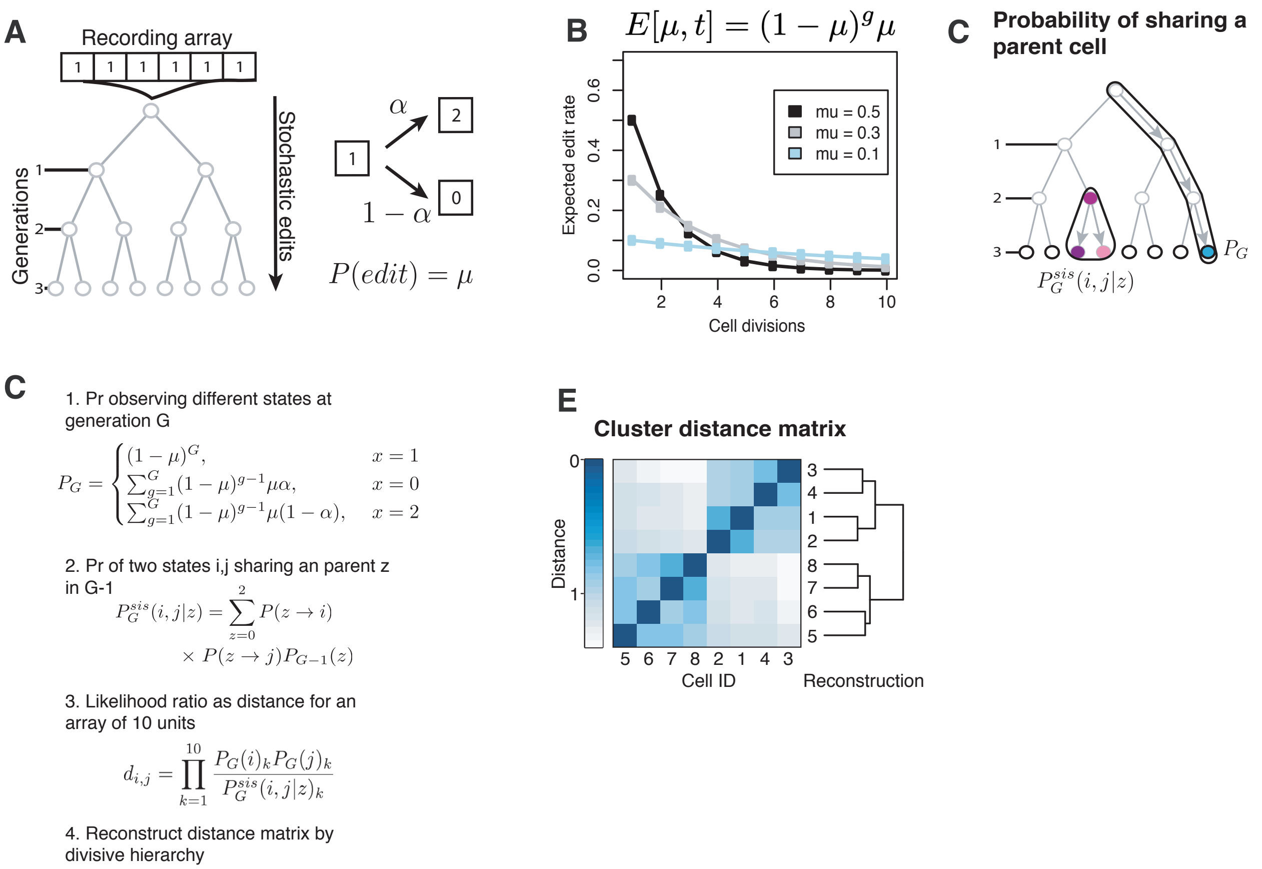


**Fig. S4. Lineage simulation and reconstruction based on maximum likelihood of sister relationships.**

(**A**) Schematic of the lineage simulation method. We define a two-parameter stochastic model where $\mu$ equals the edit rate in units of edits per site per generation, and $\alpha$ denotes the probability that the edit goes to state 2**.** The model assumes that cells divide synchronously and at a constant rate such that at generation G the lineage comprises $2^G$ cells. (**B**) For a constant $\mu$, the number of new edits appearing in each generation decays exponentially as dictated by the equation $E[\mu, t] = (1-\mu)^g\mu$ where g is the number of generations (cell divisions) and $E[\mu, t]$ is the expected fraction of edited sites. For intMEMOIR, the experimental value of $\mu$ is ~0.1-0.3 (**C**) Schematic of the reconstruction approach. We first compute the probability that a trit is in either of the three possible states at generation G, combining the transition probabilities shown in A and the equation from B and called this distribution $P_G$ (blue cell). For sister likelihood, we compute the probability that two cells $(i, j)$ share a parent $z$ in the previous generation. (**D**) Equations for reconstructing lineages based on sister likelihood. 1. The probability that a recording unit is in either of the three possible states at generation G, independently of the other cells. 2. The probability that a parent cell $z$ at $G-1$ transitions into the states i, j. This equation assumes that the daughter cells $(i, j)$ inherit the state of $z$ and then edit with probability $\mu$. Since the recording is irreversible, the only valid transitions are $1 \to 0$ and $1 \to 2$; once a cell reaches either state 2 or 0, all its daughters will inherit that state with Pr =1. We finally sum over all possible states of the parent cell $z$. 3. We can then calculate the joint probability for the 10 units as the product of the probabilities of each unit. And compare this number to the probabilities of observing the states $(i, j)$ assuming no sister relationship, which are just the product of their $P_G$ probabilities in the numerator. This ratio quantifies the likelihood of observing a given pair of array states for two sister cells compared to two unrelated cells. 4. This likelihood provides a pairwise distance metric that we then use to reconstruct the lineage tree. (**E**) Once we computed the likelihood ratio for all pairs of cells, we can cluster the matrix using divisive hierarchical clustering, which starts by partitioning the data set into the most distinct groups, then it proceeds to partition each subgroup into two groups iteratively until each group contains only one cell. Ideally, each partition of the algorithm would correspond to a cell division event.

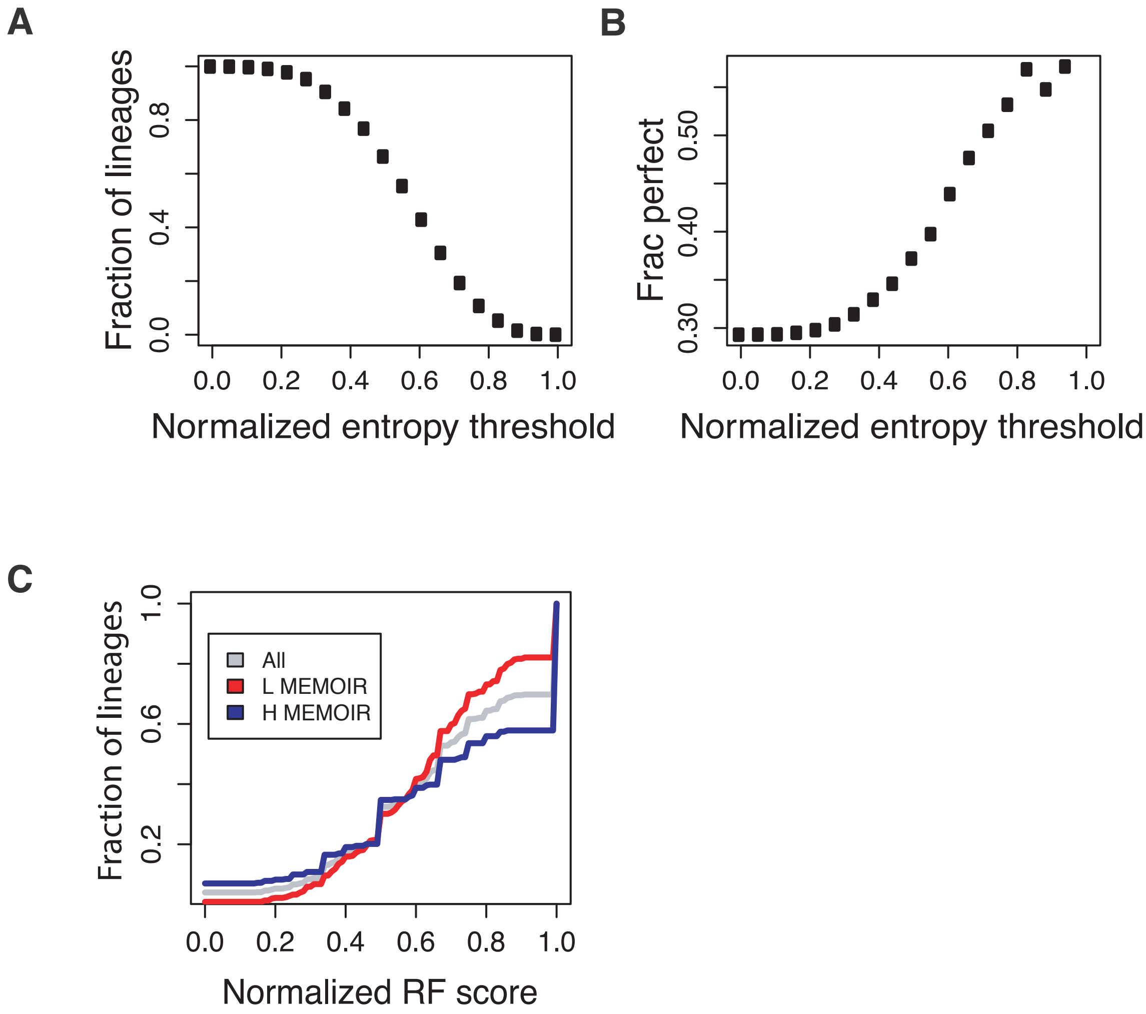


**Fig. S5. Barcode entropy enriches for colonies that reconstruct with greater accuracy.** We computed the entropy for a lineage as the sum of the individual entropies for each trit, using Shannon's formula. The normalized entropy is then computed as the lineage's entropy times the fraction of edited sites for that lineage, scaled by the maximum such that the metric has a range from [0,1]. This simulated dataset comprises 3000 lineages. (**A**) The fraction of lineages with normalized entropy larger than the threshold. (**B**) The fraction of perfectly reconstructed lineages for increasing thresholds of normalized entropy. For a given threshold value, we split the dataset and calculated the fraction of perfect trees in the high-entropy set. Note that the number of lineages analyzed decreases with increasing entropy thresholds, as shown in (A). (**C**) As an example, using a threshold of 0.6, we obtain a high-accuracy set of colonies that exhibit a fraction of perfect trees > 0.4. Note that the threshold is arbitrary and can be tuned to maximize the numbers of colonies and minimize the false discovery rate.

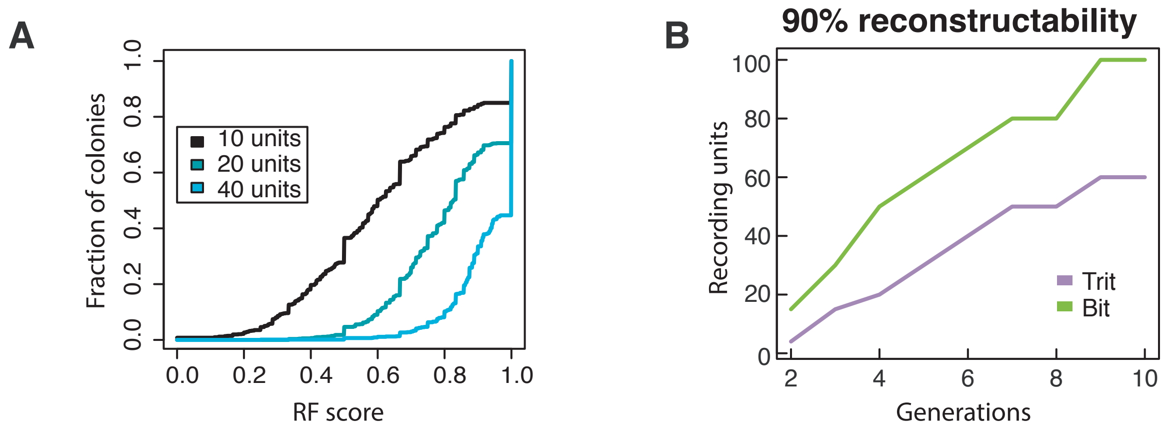


**Fig. S6. Additional intMEMOIR arrays increase reconstruction accuracy and depth.**
(**A**) Accuracy in the reconstruction of simulated lineages using increasing numbers of recording units. Parameters were estimated from experimental data. The structure of the lineage trees used in the simulation are those observed experimentally. Using 40 units arranged as 4 intMEMOIR arrays, more than 50% of lineages can be reconstructed perfectly. (**B**) For a given accuracy (90%), the number of recording units necessary for reconstruction scales with the depth of the lineage. The calculation assumes binary lineages with no cell death.

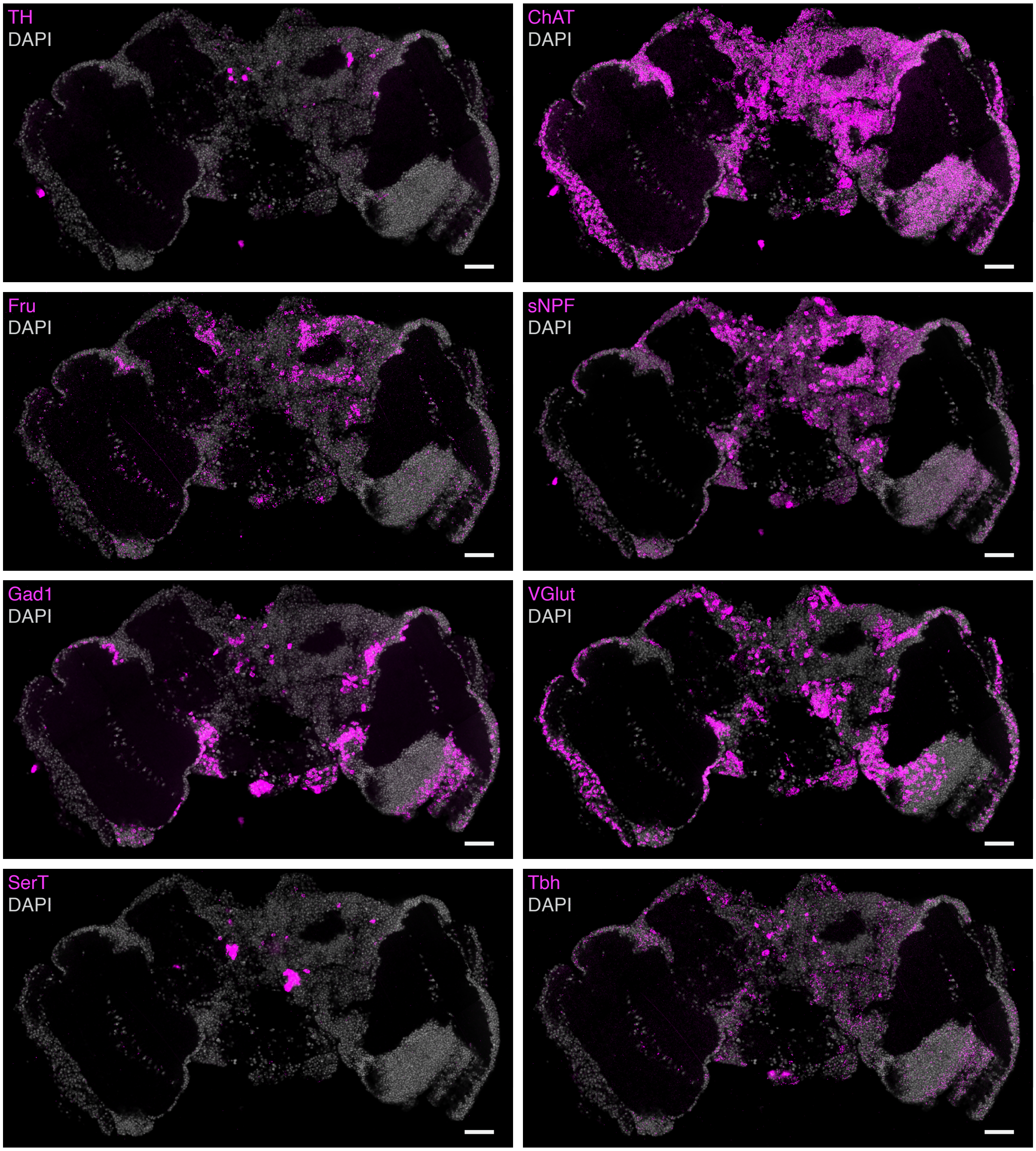


**Fig. S7. Probing the expression of 8 endogenous genes in an adult *Drosophila* brain section with smFISH.**

In addition to the intMEMOIR array, we probed for the expression of 8 endogenous genes in the same brain section: tyrosine hydroxylase (TH), choline acetyltransferase (ChAT), fruitless (Fru), short neuropeptide F precursor (sNPF), glutamic acid decarboxylase (Gad1), vesicular glutamate transporter (VGlut), serotonin transporter (SerT), and tyramine ß-hydroxylase (Tbh). Endogenous genes and DAPI signals are shown in magenta and gray, respectively (scale bar, 30 µm).

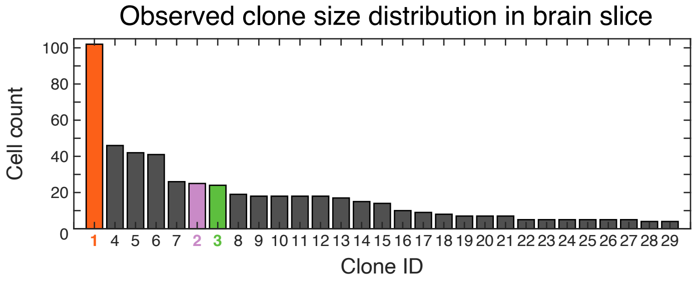


**Fig. S8. intMEMOIR recovers clone sizes *in vivo*.**

Cells in the adult *Drosophila* brain section were segmented, their array states determined (see Materials and Methods), and clones with ≥four cells and at least one unit inverted were chosen for downstream analysis. Clones 1-3, as shown in Fig. 5E, are highlighted in their corresponding colors.

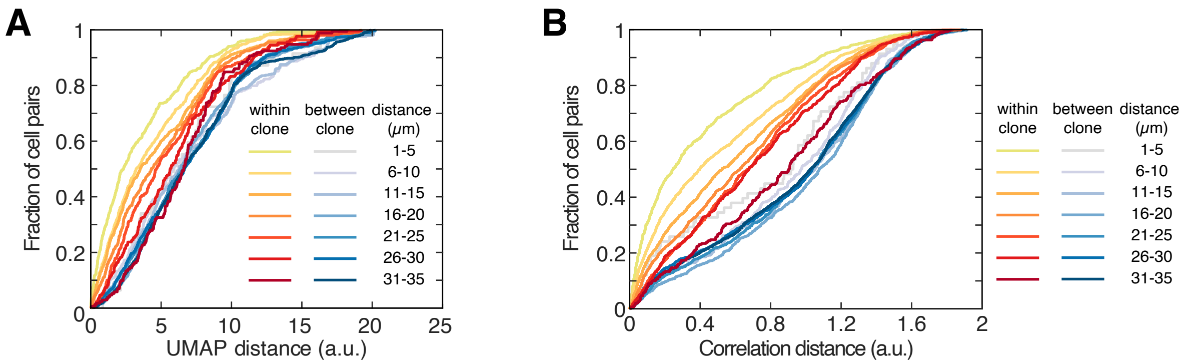


**Fig. S9. Intra-lineage spatial distribution of cells predicts fate similarity in subsamples or with alternative distance metric.**

(**A**) To determine if the relationship observed in Fig. 5M was skewed by the large Kenyon cell clone, we repeated the analysis omitting those cells. The results still showed a strong role for lineage in cell fate determination at close distances. (**B**) Clonal dependence of cell type similarity at short distances is observed when we use Pearson correlation to measure gene expression similarity, demonstrating that it is robust to the choice of distance metric.

### References


1. C. Blanpain, B. D. Simons, Unravelling stem cell dynamics by lineage tracing. *Nat. Rev. Mol. Cell Biol.* 14, 489–502 (2013).

2. K. Kretzschmar, F. M. Watt, Lineage tracing. *Cell*. 148, 33–45 (2012).

3. M. B. Woodworth, K. M. Girskis, C. A. Walsh, Building a lineage from single cells: genetic techniques for cell lineage tracking. *Nat. Rev. Genet.* 18, 230–244 (2017).

4. S.-H. S. Wu, J.-H. Lee, B.-K. Koo, Lineage Tracing: Computational Reconstruction Goes Beyond the Limit of Imaging. *Mol. Cells*. 42, 104–112 (2019).

5. C. S. Baron, A. van Oudenaarden, Unravelling cellular relationships during development and regeneration using genetic lineage tracing. *Nat. Rev. Mol. Cell Biol.* 20, 753–765 (2019).

6. D. Frumkin, A. Wasserstrom, S. Kaplan, U. Feige, E. Shapiro, Genomic variability within an organism exposes its cell lineage tree. *PLoS Comput. Biol.* 1, e50 (2005).

7. S. J. Salipante, M. S. Horwitz, Phylogenetic fate mapping. *Proc. Natl. Acad. Sci. U. S. A.* 103, 5448–5453 (2006).

8. N. Navin, J. Kendall, J. Troge, P. Andrews, L. Rodgers, J. McIndoo, K. Cook, A. Stepansky, D. Levy, D. Esposito, L. Muthuswamy, A. Krasnitz, W. R. McCombie, J. Hicks, M. Wigler, Tumour evolution inferred by single-cell sequencing. *Nature*. 472, 90–94 (2011).

9. S. Behjati, M. Huch, R. van Boxtel, W. Karthaus, D. C. Wedge, A. U. Tamuri, I. Martincorena, M. Petljak, L. B. Alexandrov, G. Gundem, P. S. Tarpey, S. Roerink, J. Blokker, M. Maddison, L. Mudie, B. Robinson, S. Nik-Zainal, P. Campbell, N. Goldman, M. van de Wetering, E. Cuppen, H. Clevers, M. R. Stratton, Genome sequencing of normal cells reveals developmental lineages and mutational processes. *Nature*. 513, 422–425 (2014).

10. M. A. Lodato, M. B. Woodworth, S. Lee, G. D. Evrony, B. K. Mehta, A. Karger, S. Lee, T. W. Chittenden, A. M. D'Gama, X. Cai, L. J. Luquette, E. Lee, P. J. Park, C. A. Walsh, Somatic mutation in single human neurons tracks developmental and transcriptional history. *Science*. 350, 94–98 (2015).

11. R. U. Sheth, H. H. Wang, DNA-based memory devices for recording cellular events. *Nat. Rev. Genet.* 19, 718–732 (2018).

12. L. S. Ludwig, C. A. Lareau, J. C. Ulirsch, E. Christian, C. Muus, L. H. Li, K. Pelka, W. Ge, Y. Oren, A. Brack, T. Law, C. Rodman, J. H. Chen, G. M. Boland, N. Hacohen, O. Rozenblatt-Rosen, M. J. Aryee, J. D. Buenrostro, A. Regev, V. G. Sankaran, Lineage Tracing in Humans Enabled by Mitochondrial Mutations and Single-Cell Genomics. *Cell*. 176, 1325–1339.e22 (2019).

13. H. Zong, J. S. Espinosa, H. H. Su, M. D. Muzumdar, L. Luo, Mosaic analysis with double markers in mice. *Cell*. 121, 479–492 (2005).

14. J. Livet, T. A. Weissman, H. Kang, R. W. Draft, J. Lu, R. A. Bennis, J. R. Sanes, J. W. Lichtman, Transgenic strategies for combinatorial expression of fluorescent proteins in the nervous system. *Nature*. 450, 56–62 (2007).

15. H.-H. Yu, C.-H. Chen, L. Shi, Y. Huang, T. Lee, Twin-spot MARCM to reveal the developmental origin and identity of neurons. *Nat. Neurosci.* 12, 947–953 (2009).

16. S. D. Perli, C. H. Cui, T. K. Lu, Continuous genetic recording with self-targeting CRISPR-Cas in human cells. *Science*. 353 (2016), doi:10.1126/science.aag0511.

17. A. McKenna, G. M. Findlay, J. A. Gagnon, M. S. Horwitz, A. F. Schier, J. Shendure, Whole-organism lineage tracing by combinatorial and cumulative genome editing. *Science*. 353, aaf7907 (2016).

18. W. Pei, T. B. Feyerabend, J. Rössler, X. Wang, D. Postrach, K. Busch, I. Rode, K. Klapproth, N. Dietlein, C. Quedenau, W. Chen, S. Sauer, S. Wolf, T. Höfer, H.-R. Rodewald, Polylox barcoding reveals haematopoietic stem cell fates realized in vivo. *Nature*. 548, 456–460 (2017).

19. R. Kalhor, K. Kalhor, L. Mejia, K. Leeper, A. Graveline, P. Mali, G. M. Church, Developmental barcoding of whole mouse via homing CRISPR. *Science*. 361 (2018), doi:10.1126/science.aat9804.

20. W. Tang, D. R. Liu, Rewritable multi-event analog recording in bacterial and mammalian cells. *Science*. 360 (2018), doi:10.1126/science.aap8992.

21. A. Alemany, M. Florescu, C. S. Baron, J. Peterson-Maduro, A. van Oudenaarden, Whole-organism clone tracing using single-cell sequencing. *Nature*. 556, 108–112 (2018).

22. M. M. Chan, Z. D. Smith, S. Grosswendt, H. Kretzmer, T. M. Norman, B. Adamson, M. Jost, J. J. Quinn, D. Yang, M. G. Jones, A. Khodaverdian, N. Yosef, A. Meissner, J. S. Weissman, Molecular recording of mammalian embryogenesis. *Nature*. 570, 77–82 (2019).

23. K. L. Frieda, J. M. Linton, S. Hormoz, J. Choi, K.-H. K. Chow, Z. S. Singer, M. W. Budde, M. B. Elowitz, L. Cai, Synthetic recording and in situ readout of lineage information in single cells. *Nature*. 541, 107–111 (2017).

24. E. Lubeck, A. F. Coskun, T. Zhiyentayev, M. Ahmad, L. Cai, Single-cell in situ RNA profiling by sequential hybridization. *Nat. Methods*. 11 (2014), pp. 360–361.

25. C.-H. L. Eng, M. Lawson, Q. Zhu, R. Dries, N. Koulena, Y. Takei, J. Yun, C. Cronin, C. Karp, G.-C. Yuan, L. Cai, Transcriptome-scale super-resolved imaging in tissues by RNA seqFISH. *Nature*. 568, 235–239 (2019).

26. A. Raj, P. van den Bogaard, S. A. Rifkin, A. van Oudenaarden, S. Tyagi, Imaging individual

mRNA molecules using multiple singly labeled probes. *Nat. Methods*. 5, 877–879 (2008).

27. I. Salvador-Martínez, M. Grillo, M. Averof, M. J. Telford, Is it possible to reconstruct an accurate cell lineage using CRISPR recorders? *Elife*. 8 (2019), doi:10.7554/eLife.40292.

28. W. M. Stark, M. R. Boocock, D. J. Sherratt, Catalysis by site-specific recombinases. *Trends Genet.* 8, 432–439 (1992).

29. M. C. A. Smith, R. Till, M. C. M. Smith, Switching the polarity of a bacteriophage integration system. *Mol. Microbiol.* 51, 1719–1728 (2004).

30. J. Bonnet, P. Subsoontorn, D. Endy, Rewritable digital data storage in live cells via engineered control of recombination directionality. *Proc. Natl. Acad. Sci. U. S. A.* 109, 8884–8889 (2012).

31. K. Rutherford, P. Yuan, K. Perry, R. Sharp, G. D. Van Duyne, Attachment site recognition and regulation of directionality by the serine integrases. *Nucleic Acids Res.* 41, 8341–8356 (2013).

32. P. C. M. Fogg, S. Colloms, S. Rosser, M. Stark, M. C. M. Smith, New applications for phage integrases. *J. Mol. Biol.* 426, 2703–2716 (2014).

33. Z. Xu, L. Thomas, B. Davies, R. Chalmers, M. Smith, W. Brown, Accuracy and efficiency define Bxb1 integrase as the best of fifteen candidate serine recombinases for the integration of DNA into the human genome. *BMC Biotechnol.* 13, 87 (2013).

34. C. A. Merrick, J. Zhao, S. J. Rosser, Serine Integrases: Advancing Synthetic Biology. *ACS Synth. Biol.* 7, 299–310 (2018).

35. B. P. Zambrowicz, A. Imamoto, S. Fiering, L. A. Herzenberg, W. G. Kerr, P. Soriano, Disruption of overlapping transcripts in the ROSA geo 26 gene trap strain leads to widespread expression of -galactosidase in mouse embryos and hematopoietic cells. *Proceedings of the National Academy of Sciences*. 94 (1997), pp. 3789–3794.

36. P. Soriano, Generalized lacZ expression with the ROSA26 Cre reporter strain. *Nat. Genet.* 21, 70–71 (1999).

37. M. Sadelain, E. P. Papapetrou, F. D. Bushman, Safe harbours for the integration of new DNA in the human genome. *Nat. Rev. Cancer*. 12, 51–58 (2011).

38. H. A. Grunwald, V. M. Gantz, G. Poplawski, X.-R. S. Xu, E. Bier, K. L. Cooper, Super-Mendelian inheritance mediated by CRISPR-Cas9 in the female mouse germline. *Nature*. 566, 105–109 (2019).

39. I. Espinosa-Medina, J. Garcia-Marques, C. Cepko, T. Lee, High-throughput dense reconstruction of cell lineages. *Open Biol.* 9, 190229 (2019).

40. P. Ghosh, A. I. Kim, G. F. Hatfull, The orientation of mycobacteriophage Bxb1 integration

is solely dependent on the central dinucleotide of attP and attB. *Mol. Cell*. 12, 1101–1111 (2003).

41. S. D. Colloms, C. A. Merrick, F. J. Olorunniji, W. M. Stark, M. C. M. Smith, A. Osbourn, J. D. Keasling, S. J. Rosser, Rapid metabolic pathway assembly and modification using serine integrase site-specific recombination. *Nucleic Acids Res.* 42, e23 (2014).

42. H. Zeng, K. Horie, L. Madisen, M. N. Pavlova, G. Gragerova, A. D. Rohde, B. A. Schimpf, Y. Liang, E. Ojala, F. Kramer, P. Roth, O. Slobodskaya, I. Dolka, E. A. Southon, L. Tessarollo, K. E. Bornfeldt, A. Gragerov, G. N. Pavlakis, G. A. Gaitanaris, An inducible and reversible mouse genetic rescue system. *PLoS Genet.* 4, e1000069 (2008).

43. M. Iwamoto, T. Björklund, C. Lundberg, D. Kirik, T. J. Wandless, A general chemical method to regulate protein stability in the mammalian central nervous system. *Chem. Biol.* 17, 981–988 (2010).

44. H. M. T. Choi, M. Schwarzkopf, M. E. Fornace, A. Acharya, G. Artavanis, J. Stegmaier, A. Cunha, N. A. Pierce, Third-generation in situ hybridization chain reaction: multiplexed, quantitative, sensitive, versatile, robust. *Development*. 145 (2018), doi:10.1242/dev.165753.

45. D. F. Robinson, L. R. Foulds, Comparison of phylogenetic trees. *Math. Biosci.* 53, 131–147 (1981).

46. K. Sugino, J. Garcia-Marques, I. Espinosa-Medina, T. Lee, Theoretical modeling on CRISPR-coded cell lineages: efficient encoding and optimal reconstruction. *bioRxiv* (2019), p. 538488.

47. J. M. Kebschull, A. M. Zador, Cellular barcoding: lineage tracing, screening and beyond. *Nat. Methods*. 15, 871–879 (2018).

48. T. Lee, Wiring the Drosophila Brain with Individually Tailored Neural Lineages. *Curr. Biol.* 27, R77–R82 (2017).

49. R. Urbach, G. M. Technau, Neuroblast formation and patterning during early brain development in Drosophila. *Bioessays*. 26, 739–751 (2004).

50. S. R. Spindler, V. Hartenstein, The Drosophila neural lineages: a model system to study brain development and circuitry. *Dev. Genes Evol.* 220, 1–10 (2010).

51. A. C. Groth, M. Fish, R. Nusse, M. P. Calos, Construction of transgenic Drosophila by using the site-specific integrase from phage phiC31. *Genetics*. 166, 1775–1782 (2004).

52. A. Akmammedov, M. Geigges, R. Paro, Single vector non-leaky gene expression system for Drosophila melanogaster. *Sci. Rep.* 7, 6899 (2017).

53. O. Riabinina, D. Luginbuhl, E. Marr, S. Liu, M. N. Wu, L. Luo, C. J. Potter, *Nat. Methods*, in press.

54. K. Dumstrei, F. Wang, C. Nassif, V. Hartenstein, Early development of the Drosophila brain: V. Pattern of postembryonic neuronal lineages expressing DE-cadherin. *J. Comp. Neurol.* 455, 451–462 (2003).

55. K. Ito, W. Awano, K. Suzuki, Y. Hiromi, D. Yamamoto, The Drosophila mushroom body is a quadruple structure of clonal units each of which contains a virtually identical set of neurones and glial cells. *Development*. 124, 761–771 (1997).

56. S.-L. Lai, T. Awasaki, K. Ito, T. Lee, Clonal analysis of Drosophila antennal lobe neurons: diverse neuronal architectures in the lateral neuroblast lineage. *Development*. 135, 2883–2893 (2008).

57. M. Ito, N. Masuda, K. Shinomiya, K. Endo, K. Ito, Systematic analysis of neural projections reveals clonal composition of the Drosophila brain. *Curr. Biol.* 23, 644–655 (2013).

58. H.-H. Yu, T. Awasaki, M. D. Schroeder, F. Long, J. S. Yang, Y. He, P. Ding, J.-C. Kao, G. Y.-Y. Wu, H. Peng, G. Myers, T. Lee, Clonal development and organization of the adult Drosophila central brain. *Curr. Biol.* 23, 633–643 (2013).

59. E. Becht, L. McInnes, J. Healy, C.-A. Dutertre, I. W. H. Kwok, L. G. Ng, F. Ginhoux, E. W. Newell, Dimensionality reduction for visualizing single-cell data using UMAP. *Nat. Biotechnol.* (2018), doi:10.1038/nbt.4314.

60. M. Ester, H.-P. Kriegel, J. Sander, X. Xu, Others, in *Kdd* (1996), vol. 96, pp. 226–231.

61. A. Askary, L. Sanchez-Guardado, J. M. Linton, D. M. Chadly, M. W. Budde, L. Cai, C. Lois, M. B. Elowitz, In situ readout of DNA barcodes and single base edits facilitated by in vitro transcription. *Nat. Biotechnol.* 38, 66–75 (2020).

62. S. Yamaguchi, Y. Kazuki, Y. Nakayama, E. Nanba, M. Oshimura, T. Ohbayashi, A method for producing transgenic cells using a multi-integrase system on a human artificial chromosome vector. *PLoS One*. 6, e17267 (2011).

63. S. Berg, D. Kutra, T. Kroeger, C. N. Straehle, B. X. Kausler, C. Haubold, M. Schiegg, J. Ales, T. Beier, M. Rudy, K. Eren, J. I. Cervantes, B. Xu, F. Beuttenmueller, A. Wolny, C. Zhang, U. Koethe, F. A. Hamprecht, A. Kreshuk, ilastik: interactive machine learning for (bio)image analysis. *Nat. Methods*. 16, 1226–1232 (2019).

64. I. T. Jolliffe, J. Cadima, Principal component analysis: a review and recent developments. *Philos. Trans. A Math. Phys. Eng. Sci.* 374, 20150202 (2016).
